# Lysosomal acid lipase regulates cholesterol metabolism during phagosomal maturation

**DOI:** 10.64898/2026.09.18.752819

**Authors:** Ojal Saharan, Bhawna, Shakuntala Surender Kumar Saraswati, Abdul Akhir, Sidharth Chopra, Dhiraj Kumar, Roop Mallik, Siddhesh S. Kamat

## Abstract

Phagocytosis, a central process in innate immunity, depends on dynamic lipid remodelling, yet how cholesterol accumulates on late phagosomes remains unresolved. Here, we identify lysosomal acid lipase (LIPA) as a key cholesterol ester (CE) hydrolase driving cholesterol mobilization during phagosomal maturation. Using integrated lipidomics, chemoproteomics, and biochemical assays, we show that LIPA exhibits acidic CE hydrolase activity enriched on late phagosomes, generating free cholesterol essential for lipid raft formation. Pharmacologically inhibiting LIPA disrupts cholesterol-rich lipid raft assembly, impairs phagosomal trafficking, and alters pathogen fate — enhancing *Staphylococcus aureus* persistence, while restricting *Mycobacterium tuberculosis* survival. These findings reveal that LIPA couples cholesterol metabolism to phagocytosis, defining a mechanistic link between lipid catabolism and antimicrobial defence. By positioning CE hydrolysis as a critical determinant of phagosomal dynamics, our work uncovers a metabolic checkpoint in innate immunity and identifies LIPA as a potential therapeutic node in infection and inflammation.

**SIGNIFICANCE STATEMENT:** Phagocytosis is a central antimicrobial process, yet the metabolic signals that enable phagosomes to mature and eliminate pathogens remain poorly defined. This study uncovers lysosomal acid lipase (LIPA) as a previously unrecognized regulator of cholesterol flux during phagosomal maturation. By integrating lipidomics, chemoproteomics, and functional biochemical assays, we show that LIPA-generated cholesterol is essential for assembling lipid raft microdomains that drive the directional trafficking of late phagosomes. Disrupting this single enzymatic step impairs phagosomal progression and alters the intracellular fate of diverse pathogens, enhancing *Staphylococcus aureus* persistence while restricting *Mycobacterium tuberculosis* survival. These findings reveal that cholesterol metabolism is not merely supportive but mechanistically integral to phagosome function. By establishing LIPA as a metabolic checkpoint that couples lipid catabolism to innate immune defence, this work highlights new opportunities to modulate host–pathogen interactions through targeted manipulation of lipid pathways.

## INTRODUCTION

Cholesterol, a fundamental structural component of cellular membranes, exerts diverse influences on immune function that extend far beyond its classical role in maintaining membrane integrity and lipid homeostasis^1,2^. Owing to its planar molecular structure, cholesterol preferentially partitions with sphingolipids to form lipid rafts^3,4^, specialized microdomains, within the plasma membrane. These cholesterol-rich rigid regions serve as organizing platforms that facilitate spatial clustering of pattern recognition receptors, such as Toll-like receptors, in macrophages and dendritic cells, thereby modulating immune signaling and pathogen recognition^5^.

Dysregulated cholesterol metabolism profoundly impacts immunity. For example, excess accumulation of free cholesterol in macrophages induces lysosomal stress and activates the inflammasome^6,7^. On the other hand, alterations in sterol biosynthesis intermediates, such as desmosterol, reprogram macrophage polarization and drastically affects innate immune memory^8,9^. Quite interestingly, impaired cellular cholesterol transport also augments proinflammatory signaling, linking sterol homeostasis directly to immune function and homeostasis. Such a metabolic–immunological crosstalk contributes to chronic inflammatory diseases like atherosclerosis and obesity, where cholesterol accumulation drives foam cell formation and systemic inflammation^10–13^. Of note, clinically relevant pathogens like *Mycobacterium tuberculosis* (*M. tuberculosis*)^14,15^, *Helicobacter pylori* (*H. pylori*)^16,17^, and *Staphylococcus aureus* (*S. aureus*)^18,19^ have evolved intricate mechanisms to evade the immune system by exploiting host cholesterol pools to establish intracellular niches^20,21^, underscoring a dual role for cholesterol in host defence and infection susceptibility.

In this context, phagocytosis is a vital immunological process, where specialized immune cells like macrophages engulf and digest pathogens, cellular debris, and foreign particles^22,23^. This evolutionary conserved process eliminates harmful microbes, presents antigens to adaptive immune cells, and releases signaling molecules that coordinate inflammation and tissue repair, maintaining host defence and systemic homeostasis. Over the past decade or so, it has been shown that cholesterol plays an indispensable role during phagosomal maturation, a critical step during phagocytosis^24^. Specifically during phagosomal maturation, relative to early phagosomes (EPs), cholesterol is significantly enriched on late phagosomes (LPs), where it partitions with ceramides (and/or glucosylceramides) to form lipid rafts, that facilitate the clustering of dynein motor proteins on LPs^25,26^. This cholesterol-dependent clustering of dyneins generates sufficient intracellular forces that drive the unidirectional motion of LPs towards the lysosomes for their eventual degradation^25^. Interestingly, in cholesterol-storage disorders, dysregulated lipid metabolism produces cholesterol-depleted phagosomes that are unable to efficiently fuse with the lysosomes^27,28^. Additionally, these cholesterol-ceramide rich lipid rafts on LPs are also important for the optimal activity of vacuolar ATPases, that are responsible for acidification of the phagosome lumen during phagosomal maturation^24,26^.

While lipidomics and cell biological studies have shown that cholesterol is enriched on LPs, mechanisms underlying as to how cholesterol accumulates on LPs remain unsolved. To address this problem, using complementary liquid chromatography coupled to mass spectrometry (LC-MS) based lipidomics platforms, biochemical assays and chemoproteomics, we identify the lysosomal acid lipase (LIPA) as a putative cholesterol ester hydrolase during phagosomal maturation. We find that inhibiting LIPA activity disrupts cholesterol-rich lipid raft formation on LPs, impairs phagosomal maturation, and has interesting effects on the intracellular persistence of *M. tuberculosis* and *S. aureus*. These findings reveal a previously unrecognized role for LIPA in linking cholesterol metabolism to phagocytosis and innate antimicrobial defence, highlighting cholesterol as a key central metabolic determinant of immune cell function.

## RESULTS

### Identification of a putative cholesterol ester hydrolase activity during phagocytosis

Recently, we developed a highly sensitive and quantitative LC-MS method to profile free cholesterol (hereafter referred to as cholesterol) and its fatty acid esters (cholesterol esters, CEs) in various mammalian cells and tissues^29^. Given the importance of cholesterol during phagocytosis, we decided to apply this LC-MS method to quantify cholesterol and CEs on purified EPs and LPs. Consistent with previous findings^25,30^, we found that relative to EPs, LPs had significantly more cholesterol (∼ 3-fold) (**Figure 1A**). Interestingly, we found that concomitant to the increased cholesterol levels, LPs had substantially lower concentrations of CEs relative to EPs (∼ 2-fold) (**Figure 1A, Supplementary Figure 1**). The counter trends for cholesterol and CEs levels on EPs and LPs led us to hypothesize that there perhaps exists a putative CE hydrolase (**Figure 1B**), whose enzymatic activity might be responsible for producing “free” cholesterol from CEs during phagocytosis.

**Figure 1.**
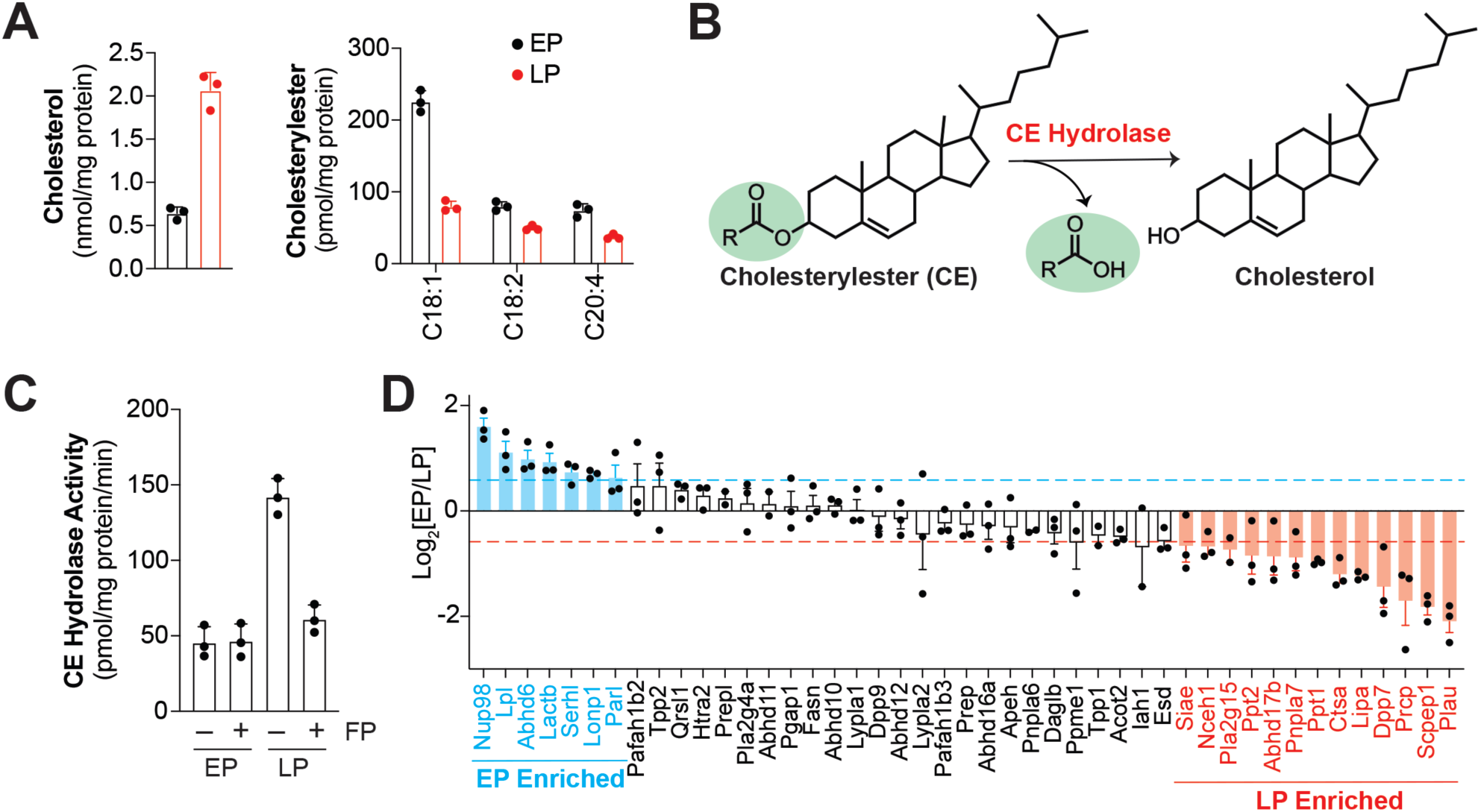
Identification of a putative CE hydrolase during phagosomal maturation. (**A**) Concentration of cholesterol and abundant CE on purified EPs (in black) and LPs (in red) measured by LC-MS analysis. (**B**) Biochemical reaction catalyzed by a putative CE hydrolase, where the CE substrate is converted to cholesterol and free fatty acid products. (**C**) CE hydrolase activity of lysates prepared from purified EPs and LPs after pretreatment with vehicle (DMSO) or a FP-probe (FP-alkyne, 50 μM, 30 min, 37 °C). (**D**) LC-MS based ABPP analysis quantitatively comparing mSH activities between proteomes prepared from purified EPs and LPs. The y-axis represents the log_2_-scale value of the enrichment ratio (EP/LP) for a particular mSH. The mSHs colored in blue and red are enriched on EPs and LPs respectively. Amongst the LP-enriched mSHs, LIPA is the most enriched neutral lipid hydrolase. For (**A**), (**C**) and (**D**), data represents mean ± standard deviation from three biological replicates per experimental group. See **Supplementary Table 1** for complete statistical analysis for various experiments described in this figure.

To test this notion, we developed a LC-MS based substrate assay to quantify the CE hydrolase activity of lysates generated from purified EPs and LPs. Of note, phagocytosis involves an intricate luminal pH gradient during phagosomal maturation, where the physiological pH of EPs and LPs is found to be approximately 7.5 and 4.5 respectively^31^. Hence, the CE hydrolase activity assays for EP and LP lysates were done at pH 7.5 and 4.5 respectively. Consistent with LPs having heightened cholesterol and lowered CEs levels relative to EPs (**Figure 1A**), we found that LP lysates possessed substantially higher acidic CE hydrolase activity (∼ 3-fold) (**Figure 1C**). Several members of the metabolic serine hydrolase (mSH) family possess lipid hydrolase (or lipase) activity against neutral lipids^32^. Hence, next, we wanted to determine if any mSH was responsible for the increased CE hydrolase activity on LPs. To confirm this, we incubated the EP and LP lysates with a fluorophosphonate (FP) probe (FP-alkyne, 50 μM, 30 min), as FP-probes are potent broad-spectrum inhibitors for the mSH enzyme class^32^. Here, we found that FP-treatment had no effect on the CE hydrolase activity of EP lysates (**Figure 1C**). However, FP-treatment completely inhibited all the increased acidic CE hydrolase activity from LP lysates, such that FP-treated LP lysates had CE hydrolase activity equivalent to EP lysates (**Figure 1C**). This result suggested that heightened acidic CE hydrolase acidic from LP lysates likely comes from a mSH that is enriched on LPs during phagosomal maturation.

Activity-based protein profiling (ABPP) has proved to be an invaluable tool in assessing mSH activities in diverse physiological settings^33^. Hence, we decided to use this chemoproteomics strategy to identify the putative mSH responsible for the heightened CE hydrolase activity on LPs. To the best our knowledge ABPP analysis have not been performed on maturing phagosomes to map mSH activities. Hence, first, we decided to perform a gel-based ABPP experiment to detect mSHs (if any) on EP/LP lysates. From this gel-based ABPP experiment, we found that both native EP/LP lysates, but not the corresponding denatured EP/LP lysates, contain numerous active mSHs (**Supplementary Figure 2**). Next, to identify these mSHs, we performed quantitative LC-MS based ABPP experiments individually on native versus denatured EP/LP lysates and found consistent with the gel-based ABPP experiment, that EP/LP lysates contained several active mSH enzymes (**Supplementary Figure 3**).

Having established the presence of active mSHs on EP/LP lysates, next, we decided to determine if a subset of these were specifically enriched on EPs or LPs. For this, we performed a quantitative LC-MS based ABPP experiment directly comparing mSH activities from equal amounts (1 mL of 1 mg/mL) of EP versus LP proteomes. For an active mSH to be considered for subsequent analysis, it needed to have ≥ 3 quantifiable peptides in ≥ 2 replicates in this quantitative LC-MS based ABPP experiment. Using these criteria, we collectively identified a total of 45 active mSHs present on EP/LP lysates (**Figure 1D**). For an active mSH to be considered enriched on EPs, it needed to have an enrichment ratio of 1.50 (0.585 on the log_2_ scale). On the other hand, an active mSH needed to have an enrichment ratio of 0.67 (1/1.5; -0.585 on the log_2_ scale) to be classified as enriched on LPs. Based on this analysis, we found that of the identified 45 active mSH, 6 and 13 were found to be enriched in the EPs and LPs proteomes respectively (**Figure 1D**). Of the 13 LP-enriched mSHs, we found a mix of proteases/peptidases (e.g. Plau, Scpep1, Prcp, Dpp7), lipases (Lipa, Pnpla7, Pla2g15, Nceh1) and small-molecule hydrolases (e.g. Ppt1/2, Siae). Based on the CE hydrolase activity data (**Figure 1C**), we specifically looked for LP-enriched lipases (acting on neutral lipids) that were mSHs. Interestingly, we found that the most enriched lipase was the lysosomal acid lipase (LIPA) (> 2-fold higher activity on LPs) (**Figure 1D**). This LP-enrichment of LIPA activity is consistent with publicly available quantitative proteomics datasets, that show heightened expression of LIPA on LPs relative to EPs (> 2-fold protein abundance on LPs)^34,35^. LIPA, a lipase from the mSH family, putatively hydrolyzes neutral lipids^32^, and given its lysosomal localization is active at an acidic pH^36–39^. Given its enrichment on LPs, and the putative enzymatic activity profile of LIPA, we decided to further characterize this lipase in the context of cholesterol metabolism during phagosomal maturation.

### LIPA functions as a cholesterol ester hydrolase during phagocytosis

The small molecule Lalistat-2 (LAL) is a selective, competitive inhibitor of mammalian LIPA (**Figure 2A**) and has potent inhibitory activity against purified human enzyme (IC_50_ ∼ 150 nM) and endogenous LIPA in different mammalian cells (IC_50_ < 1 μM)^40,41^. Therefore, we decided to use this LIPA-specific inhibitor to assess if this lysosomal lipase possesses any CE hydrolase activity during phagocytosis. First, incubated EP/LP lysates with LAL (20 μM, 30 min), and assayed them for CE hydrolase activity. We found that LAL treatment had no effect on the CE hydrolase activity from EP lysates. However, the heightened acidic CE hydrolase activity from LP lysates was completely ablated upon treatment with LAL (**Figure 2B**), suggesting that this lipase significantly contributes to the acidic CE hydrolase activity on LPs.

**Figure 2.**
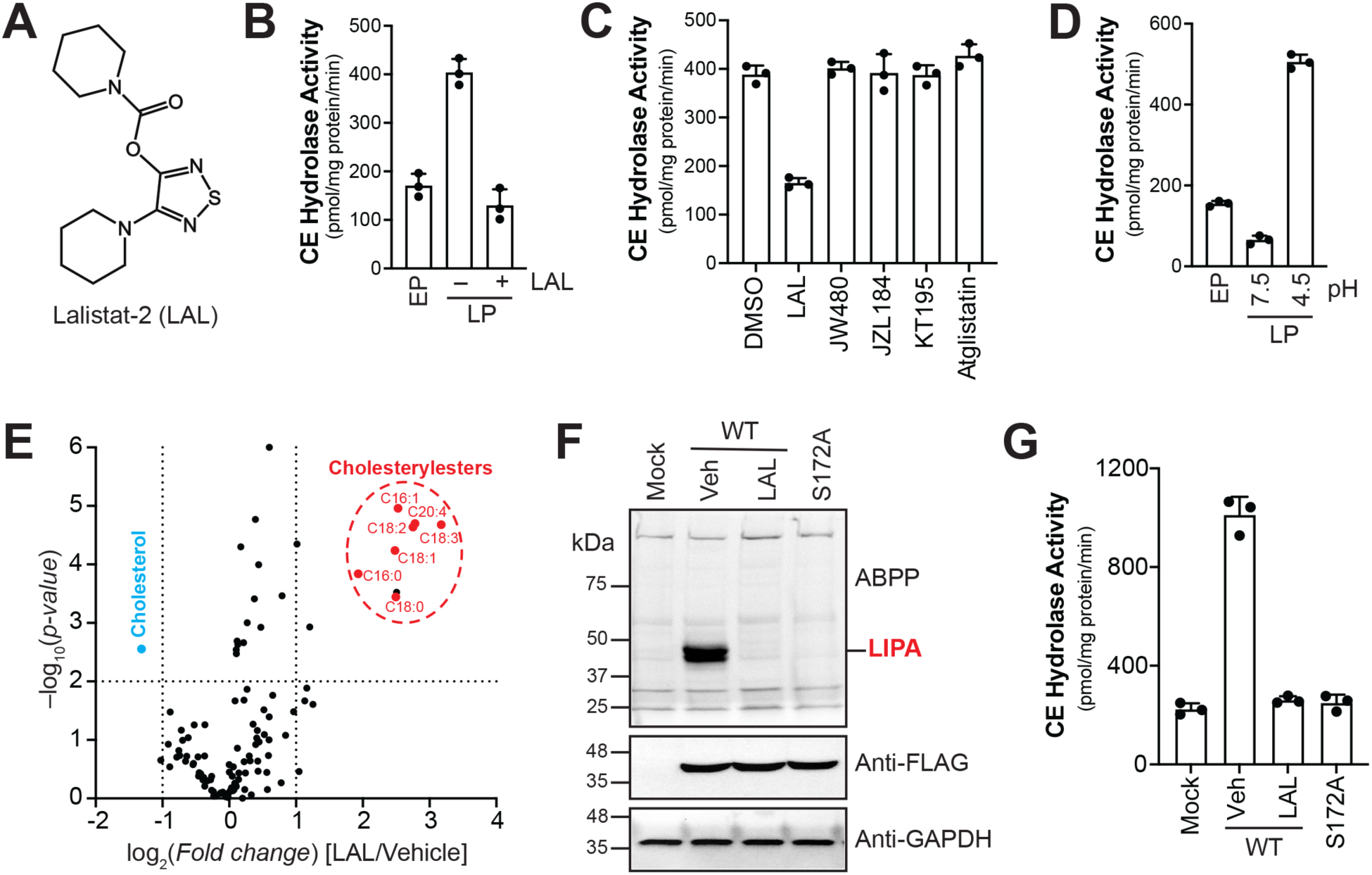
LIPA has acidic CE hydrolase activity during phagosomal maturation. (**A**) Chemical structure of Lalistat-2 (LAL), a LIPA specific reversible inhibitor. (**B**) CE hydrolase activity of lysates prepared from purified EPs and LPs after pretreatment of LP lysates with vehicle (DMSO) or LAL (20 μM, 30 min, 37 °C). (**C**) CE hydrolase activity of lysates prepared from purified LPs after pretreatment with vehicle (DMSO) or LAL or various inhibitors (20 μM, 30 min, 37 °C) that act as control for the off targets of LAL. (**D**) Comparing the CE hydrolase activity of lysates prepared from purified EPs and LPs, where the LP lysates are assayed at two pHs: acidic (pH 4.5) and physiological (pH 7.5). (**E**) Volcano plot showing differentially changing lipids on LAL-treated LPs relative to vehicle-treated LPs, as determined by LC-MS/MS analysis. Each data point represents the mean value from three independent biological replicates. The dashed line parallel to the *x*-axis denotes a cutoff with an adjusted *p*-value of 0.01, while the two dashed lines parallel to the *y*-axis denote a 2-fold change in relative lipid concentration. Based on these filtering criteria, upon LAL-treatment, CEs accumulate on LPs (shown in red), while cholesterol concomitantly decreases (shown in blue), consistent with the acidic CE hydrolase activity of LIPA. (**F**) Lysates from HEK293T cells transfected with mock (empty vector), WT mouse LIPA (vehicle (DMSO) or LAL treated (20 μM, 30 min, 37 °C)), or the S172A variant of mouse LIPA were assessed using gel-based ABPP (top panel) (to check activity of WT and S172A LIPA in HEK293T cell lysates), Western blot analysis using an anti-FLAG antibody (middle panel) (to check overexpression of WT and S172A LIPA in HEK293T cell lysates), and an anti-GAPDH antibody (bottom panel) (to confirm equal protein loading). This experiment was done 3 times with reproducible results each time. (**G**) CE hydrolase activity of HEK293T cell lysates transfected with mock (empty vector), WT mouse LIPA (vehicle (DMSO) or LAL treated (20 μM, 30 min, 37 °C)), or the S172A variant of mouse LIPA. All the assays for (**F**) and (**G**) were done at pH 4.5. For (**B**), (**C**), (**D**) and (**G**) data represents mean ± standard deviation from three biological replicates per experimental group. See **Supplementary Table 1** for complete statistical analysis for various experiments described in this figure.

LAL has the highest inhibitory activity against LIPA at an acidic pH (∼ 4.5)^40,41^. However, at higher concentrations (∼ 20 μM) and physiological pH (∼ 7.5), *in vitro* studies show that LAL also displays some inhibitory activity against a few lipases that act on neutral lipids, namely NCEH1, MAGL, ABHD6 and ATGL^40^. To rule out the possibility of any off-target effects of LAL, we assayed LP lysates for CE hydrolase activity against selective inhibitors of all the aforementioned lipases (20 μM, 30 min), i.e. JW480 for NCEH1^42^, JZL184 for MAGL^43^, KT195 for ABHD6^44^ and Atglistatin for ATGL^45^ (**Supplementary Figure 4**). From this experiment, we found that only LAL, but none of the other inhibitors, significantly reduced the acidic CE hydrolase activity from LP lysates (**Figure 2C**). LIPA is a lysosomal lipase which is most active at acidic pH (∼ 4.5) and significantly loses enzymatic activity at physiological pH (∼ 7.5)^36–39^. Consistent with this pH-activity profile of LIPA, we found that the CE hydrolase activity from LP lysates was substantially depleted when assays were done at a physiological pH (∼ 7.5) (**Figure 2D**). Thus, taken together, the results so far suggest that in the context of EPs/LPs, LAL is a specific inhibitor of LIPA, and that LIPA is indeed the lipase that is responsible for the heightened acidic CE hydrolase activity observed on LPs.

LIPA is a putative neutral lipid hydrolase, and *in vitro* substrate profiling assays show that it has enzymatic activity against an array of neutral lipids (e.g. CE, triacylglycerols, retinyl esters, diacylglycerols)^36–39^. Hence, to ascertain the lipid pathways regulated by LIPA during phagosomal maturation, we decided to purify LPs from macrophages that were treated with or without LAL and perform an untargeted lipidomics experiment on them. From this LC-MS based lipid profiling study, we found that relative to LPs purified from vehicle-treated macrophages, those purified from LAL-treated macrophages had substantially higher levels of CEs and significantly depleted cholesterol concentrations (**Figure 2E, Supplementary Table 1**). The elevation in CE levels on LPs purified from LAL-treated macrophages was seen in all the CEs detected (**Figure 2E, Supplementary Table 1**). Quite interestingly, unlike its substrate profile from *in vitro* assays, we did not see any significant change in any other neutral lipid, phospholipid or sphingolipid class between the two groups of LPs (**Supplementary Table 1**), suggesting that during phagosomal maturation LIPA specifically has acidic CE hydrolase activity.

Finally, we overexpressed the wild-type (WT) and a catalytically inactive S172A mutant of LIPA (mouse LIPA, UniProt ID: Q9Z0M5) in HEK293T cells using established transient transfection protocols^46^. By Western blot analysis, we found that upon transient transection, both WT and S172A variants of LIPA had equal overexpression in HEK293T cell lysates relative to the mock control (**Figure 2F**). Using gel-based ABPP assay, we found that consistent with the reported enzyme activity profile, WT LIPA, but not the S172A mutant of LIPA, showed robust activity at pH 4.5 relative to the mock control (**Figure 2F**). Further, we also found that treating WT LIPA with LAL (20 μM, 30 min) completely abolished all its activity in the gel-based ABPP assay (**Figure 2F**). Next, we assayed these lysates for acidic CE hydrolase activity using the LC-MS based substrate assay. Consistent with the results from the gel-based ABPP assays, we find that relative to the mock control, WT LIPA, but not the S172A mutant or LAL-treated WT LIPA, has substantial CE hydrolase activity at pH 4.5 (**Figure 2G**). Quite interestingly, we find that WT LIPA does not show any appreciable activity at pH 7.5 in both the gel-based ABPP and LC-MS based CE hydrolase assay (**Supplementary Figure 5**). Taken together, this data conclusively shows that LIPA is indeed CE hydrolase with optimal activity at acidic pH, and LAL is a potent inhibitor of this lipase.

### Disruption of LIPA activity hampers phagosomal maturation

Pharmacological inhibition of LIPA activity by LAL results in dysregulated cholesterol metabolism during phagocytosis. Hence, we wanted to see if inhibiting this acidic CE hydrolase has any effect on phagosomal maturation. Towards this, first, we checked if inhibiting LIPA affected the engulfment step by macrophages during phagocytosis. To assess this, macrophages were treated with LAL, and bead uptake (using fluorescent beads) was measured by established cellular immunofluorescence assays (IFA)^25^. From this experiment, we found that there was no change in the total number of beads taken up by macrophages when they were treated with LAL, relative to a vehicle control (**Figure 3A**). This suggested that pharmacological inhibition of the LIPA activity has no effect on the initial engulfment step during phagocytosis.

**Figure 3.**
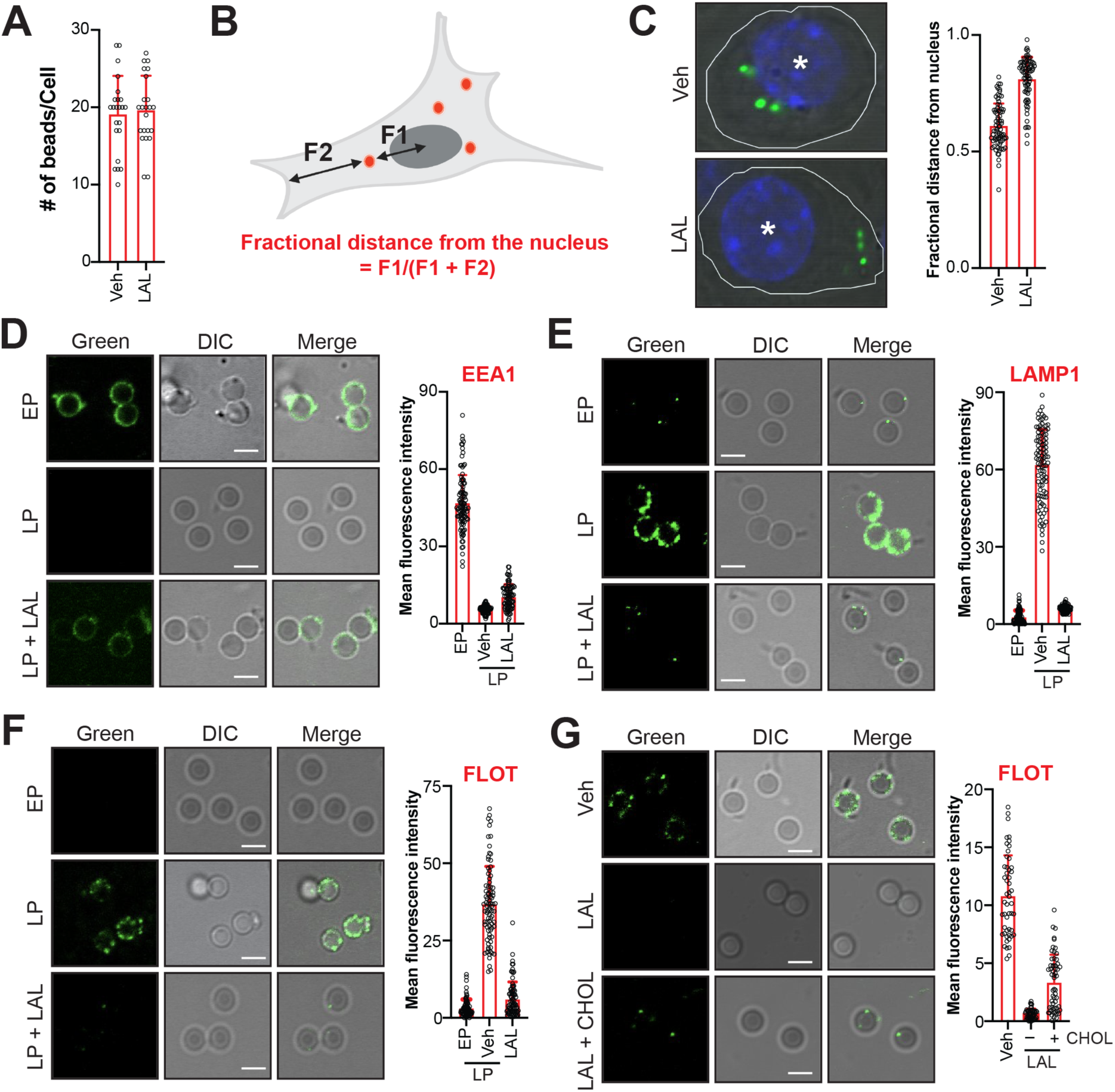
Disruption of LIPA activity affects phagosomal maturation. (**A**) Number of fluorescent beads phagocytosed by RAW264.7 cells treated with vehicle (DMSO) or LAL (20 μM, 4 hours). Data represents mean ± standard deviation from quantification from >20 cells in this experiment. (**B**) Schematic representation showing the definition of the distance of a phagocytosed bead from the center of the nucleus (F1) and from the periphery of the cell (F2). (**C**) *Left,* Representative image of fluorescent beads phagocytosed by RAW264.7 cells treated with vehicle (DMSO) or LAL (20 μM, 4 hours). *Right*, The fractional distance of phagocytosed beads from the nucleus. Data represents mean ± standard deviation from the quantification of >75 individual fluorescent beads per experimental group >20 cells. (**D-F**) *Left*, Representative images from the immunofluorescence analysis of purified EPs, LPs and LAL-treated LPs, using (**D**) anti-EEA1 antibody (EP marker), (**E**) anti-LAMP1 antibody (LP marker) and (**F**) anti-Flotilin-1 (FLOT) antibody (cholesterol-rich lipid raft marker). *Right*, Fluorescence intensity of individual phagosomes measured from the respective immunofluorescence experiment. (**G**) *Left*, Representative images from immunofluorescence analysis of LPs purified from RAW264.7 cells that were treated with vehicle, LAL (20 μM, 4 hours) and LAL (20 μM, 4 hours) supplemented with cholesterol (100 μM) using anti-FLOT antibody (cholesterol-rich lipid raft marker). *Right*, Fluorescence intensity of individual phagosomes measured in this immunofluorescence experiment. For (**D-G**) The three images per experimental panel are a green immunofluorescence channel (Green) image (for a specific protein marker), a differential interference contrast (DIC) image (for outlining the phagosome), and an overlay of the Green and DIC images (Merge). The white line on all the DIC images is a 3-μm scale. For (**D-G**) Data represents mean ± standard deviation from quantification of fluorescence intensity of >80 individual purified phagosomes per experimental group. (**A-G**) All experiments were done at least three individual times with reproducible results each time. See **Supplementary Table 1** for complete statistical analysis for various experiments described in this figure.

Another important metric in assessing the progression of phagocytosis is measuring the cellular localization of the engulfed beads from the center of the nucleus. Under normal conditions, lysosomes have a peri-nuclear localization. Hence, if phagocytosis (or phagosomal maturation) is hampered under specific conditions, the engulfed beads may not reach their eventual destination, i.e. the peri-nuclear lysosomes. This would result in an altered cellular localization of the beads taken up by the macrophages. The localization of engulfed beads (using fluorescent beads) can be quantified by measuring the “fractional distance from the nucleus” by established cellular IFA^25^ (**Figure 3B**). Quite interestingly, we found that despite having equal bead uptake, LAL-treated macrophages showed significantly higher fractional distance from the nucleus for the beads, when compared to vehicle treated macrophages (**Figure 3C**). This data suggests that the beads taken up by macrophages cannot efficiently reach the peri-nuclear lysosomes when the LIPA-dependent CE hydrolase activity is blocked.

The proteomic content of ageing phagosomes, especially EPs and LPs has been quite extensively studied^34,35,47^. As a result, various stages of phagocytosis have established proteins that serve as valuable markers to assess the progression of phagosomal maturation. Specifically, the proteins, early endosome antigen 1 (EEA1) and the lysosome-associated membrane protein 1 (LAMP1) are well-established markers of EPs and LPs respectively^22,23^. Since the disruption of LIPA activity presumably affects phagosomal maturation, next, we specifically looked for the levels of EEA1 and LAMP1 on LPs that were purified from macrophages treated with LAL. From this experiment, we found that relative to EP and LP controls, LPs purified from LAL-treated macrophages had significantly lower levels of both EEA1 (**Figure 3D**) and LAMP1 (**Figure 3E**) respectively. This suggests that while the LPs purified from LAL-treated macrophages have progressed beyond the EP-stage (reduced levels of EEA1), they have not matured to form LPs (reduced levels of LAMP1), consistent with the premise that inhibition of LIPA hampers and/or significantly delays phagosomal maturation.

Another hallmark of LPs is the formation of punctate cholesterol-rich lipid microdomains (or lipid rafts), that are essential for the clustering of dynein motor proteins, which facilitate the unidirectional transport of phagosomes to the lysosome^25^. Since LIPA functions as a CE hydrolase during phagocytosis, and its activity is essential for phagosomal maturation, we wanted to assess the extent of formation of the punctate cholesterol-rich lipid microdomains on LPs purified from LAL-treated macrophages. LPs purified from LAL-treated macrophages had substantially fewer punctate cholesterol-rich lipid microdomains compared to LPs purified from vehicle-treated macrophages (**Figure 3F**), supporting the impaired phagosomal maturation phenotype in presence of LAL. We also purified LPs from macrophages which were treated with JW480, JZL184, KT195 or Atglitatin. We found that inhibiting NCEH1, MAGL, ABHD6 or ATGL has no effect on the formation of punctate cholesterol-rich lipid microdomains on LPs (**Supplementary Figure 6**), confirming that the effects on LAL on this phenotype were indeed LIPA-specific.

Finally, to assess if exogenously supplied “free” cholesterol can rescue this phenotype and enable the formation of punctate cholesterol-rich lipid microdomains on LPs, we supplemented LAL-treated macrophages with surplus cholesterol (100 μM final concentration), and purified LPs from this condition. Quite interestingly, we found that relative to LPs purified from LAL-treated macrophages, the LPs purified from the same condition but supplemented with cholesterol had an appreciable increase (∼ 5-fold) in the punctate cholesterol-rich lipid microdomains (**Figure 3G**). However, these increased punctate cholesterol-rich lipid microdomains on LPs purified from LAL-treated macrophages supplemented with cholesterol was only about 30% of the expected amount on control LPs (**Figure 3G**). This experiment therefore suggests that exogenously supplied cholesterol can at best partially rescue this phenotype on LPs devoid of LIPA activity. Taken together, it is important to note that during phagocytosis, the *in-situ* CE hydrolase activity of LIPA produces majority of cholesterol on demand, needed to form these punctate cholesterol-rich lipid microdomains. Hence, in the absence of the enzymatic activity of LIPA (the major acidic CE hydrolase), phagosomal maturation, and in turn, the formation of the punctate cholesterol-rich lipid microdomains on LPs is considerably compromised.

### Disruption of LIPA activity affects microbial clearance via phagocytosis

The CE hydrolase activity of LIPA is critical for phagosomal maturation. However, all the assays done thus far, were performed by feeding beads to macrophages, and assessing phagosomal maturation as a function of LIPA activity. While this established strategy works as an excellent surrogate for phagocytosis, we wanted to determine the role that LIPA plays in more physiological settings during phagocytosis. To test this, we chose to perform a macrophage-based microbial clearance assay (**Figure 4A**) using two clinically relevant bacteria, namely *S. aureus* and *M. tuberculosis*. Before initiating these microbial clearance assays, we tested LAL (20 μM, 4 h) individually on both the bacterial inoculums and macrophages chosen for these experiments. We found that at the concentration and duration of testing, LAL did not have any bactericidal (**Supplementary Figure 7**) or cell death activity against macrophages.

**Figure 4.**
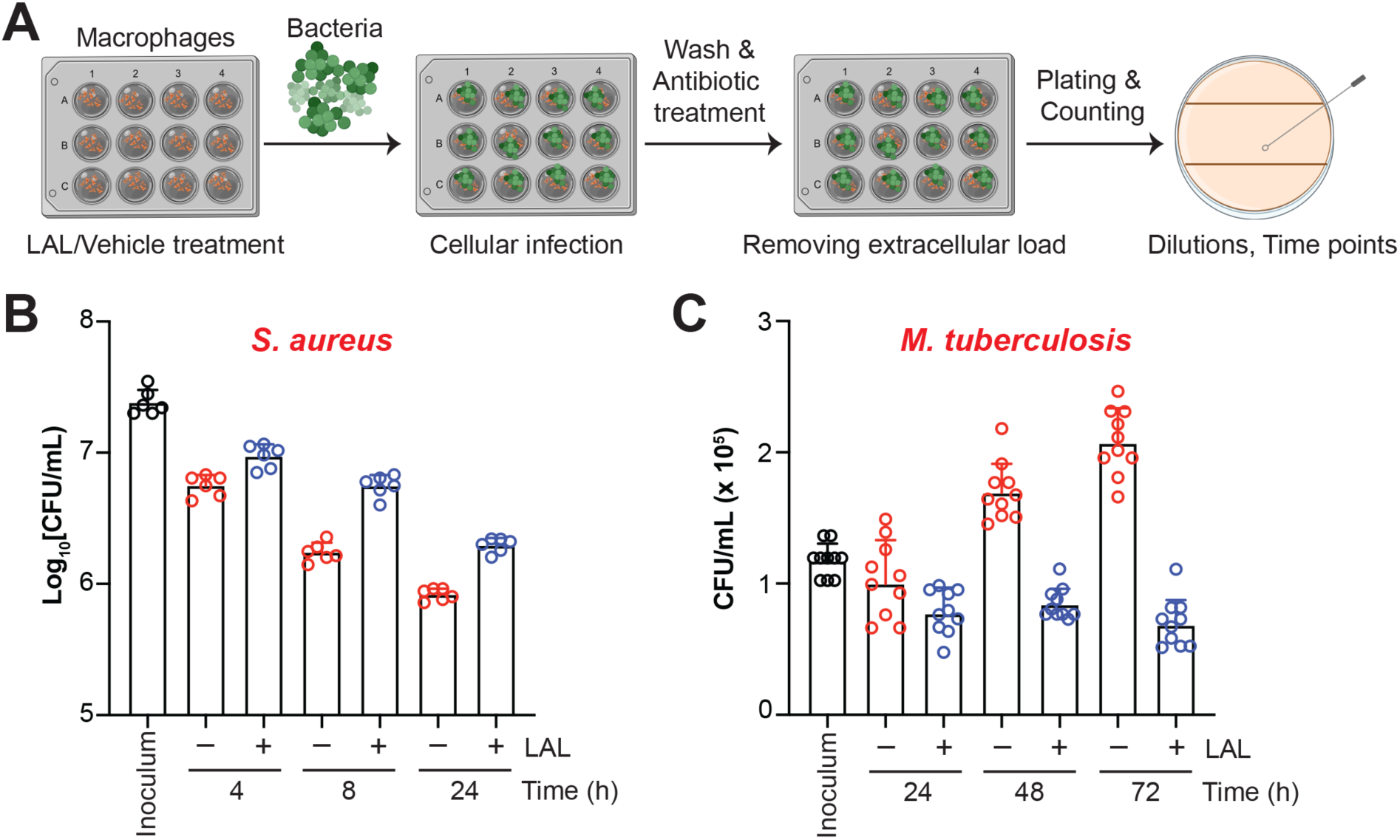
Effect of inhibiting LIPA on microbial clearance by macrophages. (**A**) Schematic representation of the microbial clearance assays. (**B, C**) Survival of (**B**) *S. aureus* and (**C**) *M. tuberculosis* in macrophages relative to the inoculum control (in black) at various time points after vehicle (in blue) or LAL treatment (20 μM) (in red). Data represents mean ± standard deviation from six (**B**) and ten (**C**) biological replicates per experimental group. All experiments were done at least three individual times with reproducible results each time. See **Supplementary Table 1** for complete statistical analysis for various experiments described in this figure.

*S. aureus* is a gram-positive bacterium with emerging resistance to known antibiotics^48,49^. It causes opportunistic skin and respiratory infections in human subjects with compromised immune systems, especially in hospital settings. However, in healthy humans, *S. aureus* is cleared by the immune system via phagocytosis. Hence, we hypothesized that if the pharmacological inhibition of LIPA hampers phagosomal maturation, there would be an increased survival of *S. aureus* in this microbial clearance assay. From this experiment, we observed that relative to the inoculum control, as a function of time, *S. aureus* survival was significantly diminished in macrophages that were vehicle-treated (**Figure 4B**). On the other hand, when phagosomal maturation is hampered by disrupting the acidic CE hydrolase activity of LIPA (LAL-treatment of macrophages), we found that substantially more *S. aureus* survived in this condition as a function of time relative to the corresponding vehicle-control (**Figure 4B**). This experiment thus suggests that *in vivo*, the CE hydrolase activity of LIPA is critical for efficient phagosomal maturation and clearance.

*M. tuberculosis* is the bacteria responsible for causing Tuberculosis, a respiratory infection that globally infects >10 million people and is responsible for >1 million deaths annually^50^. Unlike *S. aureus*, *M. tuberculosis* is resistant to phagocytic clearance and can survive (and even multiply) in macrophages^14,15^. However, *M. tuberculosis* cannot synthesize cholesterol and must therefore, scavenge cholesterol from phagosomes for its survival^21,51^. Hence, we hypothesized that *M. tuberculosis* survival can be reduced if cholesterol metabolism during phagocytosis is disturbed by inhibiting the enzymatic activity of LIPA. From this experiment, we found that relative to the inoculum control, as a function of time, *M. tuberculosis* survival increased in macrophages that were vehicle-treated (**Figure 4C**). Counter to this, we found that when the acidic CE hydrolase activity of LIPA was blocked by LAL-treatment of macrophages, the *M. tuberculosis* survival in this condition as a function of time was massively reduced relative to the corresponding vehicle-control (**Figure 4C**). Corroborating available literature, our data also shows that hijacking cholesterol during phagocytosis is critical for the survival of *M. tuberculosis* in macrophages. Interestingly, we find that the LIPA-dependent CE hydrolase activity that produces cholesterol on demand during phagosomal maturation is needed for *M. tuberculosis* survival in macrophages and shows that this cholesterol metabolism during phagocytosis has important functions in microbial clearance and survival.

## DISCUSSION

With the emergence of diverse infectious and inflammatory diseases, exploring new dimensions of innate immunity has become increasingly essential. Cholesterol, a fundamental structural constituent of cellular membranes, has long been recognized for its role in maintaining membrane integrity and fluidity^3,4^. Beyond this classical function, cholesterol has profound regulatory effects on immune signaling and pathogen clearance^1,2^. Yet, the precise biochemical mechanisms through which cholesterol modulates specific innate immune processes remain incompletely understood.

Phagocytosis, a conserved effector arm of innate immunity, represents a highly orchestrated process through which specialized phagocytes like macrophages internalize and degrade invading pathogens^22,23^. Following engulfment, nascent phagosomes undergo a sequence of maturation steps, that include the formation of EPs and LPs, of which, the latter ultimately fuses with the lysosomes leading to degradation of internalized material. This maturation process is profoundly influenced by dynamic changes in cholesterol distribution on the phagosomes^24^. Thus, understanding how cholesterol is recruited, mobilized, and regulated on phagosomes would provide critical insight into the intersection of lipid metabolism and immune regulation.

To delineate this, using integrated LC-MS based lipidomics, biochemical assays, and chemoproteomics approaches, we identify LIPA^36–39^ as a key enzyme associated with phagosomes, revealing an unrecognized role in regulating cholesterol flux during phagosomal maturation. LIPA, a CE hydrolase optimally active at acidic pH (∼4.5), exhibits markedly higher activity on LPs compared to EPs. Pharmacological inhibition of the CE hydrolase activity of LIPA by LAL disrupts this temporal lipid remodelling, resulting in impaired phagosomal maturation. This is evidenced by aberrant levels of stage-specific protein markers and hampered formation of the cholesterol-enriched lipid rafts on LPs. Notably, exogenous cholesterol supplementation fails to fully restore normal cholesterol levels in LIPA-inhibited LPs, underscoring that the on demand acidic CE hydrolase activity of LIPA, rather than simple lipid availability, governs the homeostatic balance of cholesterol during phagocytosis.

The mechanistic relevance of LIPA activity extends beyond lipid turnover to host–pathogen interactions. Microbial clearance assays reveal that the enzymatic activity of LIPA directly influences the intracellular persistence of clinically relevant pathogens such as *M. tuberculosis* and *S. aureus*. Given that cholesterol-rich lipid rafts are essential for recruiting dyneins to LPs, enabling their directional movement toward lysosomes, the CE hydrolase activity of LIPA appears to function upstream in this cholesterol-dependent trafficking pathway. By modulating cholesterol availability, the on demand activity of LIPA determines whether internalized microorganisms are successfully delivered to degradative compartments or evade destruction by residing in stalled phagosomes.

The clinical implications of this pathway are substantial. *M. tuberculosis*, the etiological agent of tuberculosis and a pathogen infecting >10 million people annually, is known to exploit host cholesterol for intracellular survival and latency^50^. Our findings demonstrate that inhibition of LIPA compromises *M. tuberculosis* persistence within phagocytes, suggesting that targeted modulation of LIPA activity may enhance host control of infection. However, this effect is context dependent. In contrast to *M. tuberculosis*, pathogens like *S. aureus* that do not rely directly on cholesterol for survival exhibit increased intracellular retention following LIPA inhibition, leading to delayed bacterial clearance. These results highlight the delicate balance required in tuning the CE hydrolase activity of LIPA, too little may hinder phagosomal progression, while excessive modulation could perturb host lipid metabolism or immune signaling. Hence, achieving an optimal balance may be imperative to leveraging LIPA as an attractive therapeutic target for treating infectious and inflammatory diseases.

A particularly intriguing question arising from our findings concerns the subcellular localization of LIPA. Conventionally regarded as a luminally oriented lipase restricted to lysosomes, LIPA’s association with phagosomes suggests a more dynamic trafficking mechanism, akin to the “kiss and run” mechanism in the biogenesis of the phagolysosome^52^. Considering the extensive vesicular communication between endosomes and lysosomes during phagocytosis, it is plausible that LIPA reaches the phagosome through transient fusion or vesicle-mediated exchange^52^. Elucidating this trafficking route could potentially uncover fundamental principles governing the redistribution of lysosomal enzymes during immune activation. Moreover, identifying the cholesterol-binding or scaffold proteins that mediate the recruitment of dyneins, may also reveal new signaling intermediates that coordinate LP trafficking and degradation during phagocytosis.

Together, these findings establish a critical mechanistic link between cholesterol metabolism and innate immune function. By identifying LIPA as a pivotal regulator of cholesterol flux during phagosome maturation, this study expands the conceptual framework of lipid–immune cross-talk. Our integrative approach of combining lipidomics, chemoproteomics, and biochemical analyses, demonstrates how a canonical degradative enzyme can dynamically influence immune processes that extend far beyond lysosomal catabolism. Interestingly, corroborating our findings, while its known that mutations to LIPA results in Wolman disease and CE storage disease (CESD) in humans^36,37^, recent clinical reports suggest that human subjects suffering Wolman disease are particularly susceptible to infections from severe inflammation ^53–55^. Additionally, in tune with recent literature, our studies support the fact that cholesterol is not merely as a structural lipid, rather an important determinant of immune efficacy, shaping the fate of pathogen clearance through spatial control of membrane dynamics^21^.

From a translational perspective, the ability to fine-tune LIPA activity presents promising opportunities for therapeutic intervention. Modulating LIPA could enhance host defence against intracellular pathogens such as *M. tuberculosis*, or conversely, dampen excessive inflammation in lipid-driven disorders where phagocytosis is compromised. By positioning cholesterol metabolism as a central regulator of immune homeostasis, this work underscores the growing convergence between metabolic and immune biology. Ultimately, the discovery of LIPA’s role in phagosomal cholesterol regulation provides a foundation for future strategies that exploit lipid metabolism to restore or enhance immune function in infectious and inflammatory diseases.

## METHODS

### Materials

Unless otherwise mentioned, all chemicals and reagents were purchased from Sigma-Aldrich, and all tissue culture media and consumables were purchased from HiMedia.

### Preparation of EPs and LPs

EPs and LPs were prepared from RAW264.7 macrophage cell line [cultured in DMEM (HiMedia; catalog # AL139A) containing 10% (v/v) fetal bovine serum (HiMedia; catalog # RM1112) and 1% (v/v) Penicillin-Streptomycin (HiMedia; catalog # A100A) at 37 °C and 5% (v/v) CO_2_] using 1-µm (Polysciences Inc.; catalog # 24326-15) or 3-µm (Polysciences Inc.; catalog # 24330-15) silica beads by a pulse chase technique previously reported by us^25,26,30^. The EP and LP preparations using 3-µm silica beads were used for the IFA assays, while those using the 1-µm silica beads were used for all the other assays. For generating inhibitor treated LPs, RAW264.7 cells were pre-treated with various inhibitors used in this study [LAL (Sigma-Aldrich; catalog # SML2053), JW480 (Sigma-Aldrich; catalog # 420175), JZL184 (Sigma-Aldrich; catalog # 475741), KT195 (Sigma-Aldrich; catalog # SML1308) or Atglistatin (Sigma-Aldrich; catalog # SML1075)] at 20 μM final concentration for 4 hours, following which LPs were prepared using protocols previously reported by us^25,26,30^.

### LC-MS method for quantifying cholesterol and CEs

The absolute concentrations of cholesterol and CEs on EPs and LPs were measured using a LC-MS recently reported by us^29^. Briefly, the protein concentrations of EP or LP preparations (prepared using 1-µm silica beads) were measured by Bradford’s reagent and adjusted to 0.5 mg/mL using sterile Dulbecco’s phosphate buffered saline (DBPS). 1 mL of this EP or LP preparation (containing 500 μg protein in total) was subjected to lipid extraction and LC-MS analysis as previously reported by us^29^. The absolute concentrations of cholesterol and CEs on EPs or LPs was calculated by measuring the area under the curve for a particular sterol species relative to the respective internal standard and then normalising this to the total protein content. In this lipid measurement experiment, cholesterol-d7 (Sigma-Aldrich; catalog # 700041P, 2 nmol per sample) and C17:0 CE (Sigma-Aldrich; catalog # 700186M, 1 nmol per sample) were used as internal standards for cholesterol and CEs respectively.

### LC-MS based CE hydrolase assay

The CE hydrolase activity was measured by a LC-MS based assay assessing free fatty acid (FFA) release from a CE substrate. The CE substrate was prepared as a 4:1 (mol%) liposomal mixture of C18:0/18:0 phosphatidylcholine (PC) (Sigma-Adrich; catalog # 850365) and C18:1 CE (Sigma-Adrich; catalog # 700269) in sodium acetate buffer (pH 4.5, 50 mM) or DPBS (pH 7.5). All assays were done in 100 μL volume, and contained protein lysates (40 μg protein), 10 μM sodium taurocholate and 100 μM C18:1 CE in the liposomal mixture. The reaction was allowed to proceed for 60 min at 37 °C with constant shaking, and subsequently quenched using 300 μL of 2:1 (v/v) chloroform (CHCl_3_): methanol (MeOH) containing 0.5 nmol of pentadecanoic acid (C15:0 FFA; Sigma-Aldrich, catalog # P6125) as an internal standard. The samples were vigorously vortexed, and centrifuged at 1500*g* for 10 min to separate organic (bottom) and aqueous (top) layers. The organic layer (bottom) was collected, and dried under a stream of nitrogen gas. The dried extracts were then reconstituted in 150 μL 2:1 (v/v) CHCl_3_:MeOH and 20 μL of this was used for the LC-MS analysis on an Agilent G6125B single-quadrupole LC-MS with electrospray ionization in the negative ion mode. The liquid chromatography (LC) solvent systems, columns and MS conditions were identical to those previously reported by us^56^. A typical LC-MS run was 21 min, starting with 0.3 mL/min 100% buffer A for 5 min, 0.5 mL/min linear gradient to 100% buffer B over 9 min, 0.5 mL/min 100% buffer B for 4 min, and equilibration with 0.5 mL/min 100% buffer A for 3 min. The product release was quantified by measuring the area under the curve for the peak corresponding to C18:1 FFA (produced from C18:1 CE) and normalizing it to the internal standard (C15:0 FFA). The substrate hydrolysis rate was corrected by subtracting the nonenzymatic rate of hydrolysis, which was obtained by using heat-denatured (15 min at 95 °C) protein lysates as reported earlier^56^.

### ABPP analysis of EPs and LPs

All gel based ABPP assays were performed using protocols previously reported by us^56,57^. Briefly, in a 50 μL reaction volume, 50 μg of protein lysate (in DPBS at pH 7.5, or in 50 mM sodium acetate buffer at pH 4.5) was incubated with 2 μM FP-rhodamine for 45 min at 37 °C with constant shaking. The reactions were quenched by adding 20 μL of 4x SDS-PAGE loading buffer followed by boiling for 5 min, and depending on the experiment, 20 or 40 μL of this sample was loaded on a 12.5% SDS-PAGE gel. Competitive gel-based ABPP experiments with LAL were done as reported earlier^46^. All gels were visualized using an iBright1500 gel documentation system (Invitrogen). All LC-MS based ABPP assays were performed using established protocols previously reported by us^46,57^. In these experiments, in a reaction volume of 1 mL, 1 mg of EP or LP proteome (in DPBS) was incubated with 50 μM FP-biotin for 60 min at 37 °C with constant shaking. Thereafter, the samples were denatured, reductively alkylated using iodoacetamide and biotinylated proteins (mSH enzymes) were enriched using avidin-chromatography. Peptides were generated by on-bead digestion using MS grade trypsin (Promega; catalog # V5111). For the downstream quantitative proteomics analysis, these tryptic peptides were reductively dimethylated (ReDiMe) using light/heavy formaldehyde using established protocols previously reported by us^46,58^. Post-labelling, the light/heavy peptides were pooled, desalted and run on a TripleTOF 6600 (Sciex) mass spectrometer fitted with an Eksigent 425 nano-LC in an information-dependent acquisition mode using LC columns, solvents and other instrumental acquisition parameters previously reported by us^59^. Peptide identification and quantification was carried out using the Protein Pilot software (version 2.0.1, Sciex) with the in-built Pro-Group and Paragon algorithms against the UniProt generated RefSeq protein database of *Mus musculus* (Release 109, last modified on 22^nd^ September 2020) as per procedures recently reported by us^59^. Denatured proteomes were prepared by boiling EP or LP lysates at 95 °C for 15 min (x3 cycles). In the assessment of mSHs activities from the native vs denatured experiments with EP/LP proteomes (**Supplementary Figure 3**), the tryptic peptides from the native and denatured proteomes were labelled with heavy and light formaldehyde respectively. In the quantification of mSH activities on EP vs LP proteomes (**Figure 1D**), the tryptic peptides from the EP and LP proteomes were labelled with heavy and light formaldehyde respectively. In this study, a maximal cut-off of 10 was imposed on the enrichment ratio (heavy label to light label) for the LC-MS based ABPP experiments.

### Untargeted lipidomics

LPs and LAL-treated LPs were prepared as mentioned earlier (prepared using 1-µm silica beads), and the lipids were enriched from these preparations (containing 500 μg protein in total) using extraction protocols previously reported by us^30^. The lipid extracts were reconstituted in 200 μL CHCl_3_:MeOH, and 10 μL of this was injected into an Agilent 6545 quadrupole time-of-flight (QTOF) LC-MS system in high-resolution auto MS/MS mode for semiquantitative untargeted lipid profiling using the electrospray ionization technique. The LC solvent system, solvent gradient, column and MS parameters were identical to those previously reported by us^56^. For lipid identification, a curated personal compound database library was curated and used. Peaks were validated based on accurate mass (± 5 ppm error), relative retention time, and MS/MS fragmentation patterns. Quantification of individual lipid species was carried out by calculating the peak areas, normalizing them to the respective internal standard spiked into the samples, and to the total protein content of the sample.

### Overexpression of LIPA

The cDNA of full-length WT LIPA (mouse, UniProt ID: Q9Z0M5) was amplified from an inhouse cDNA library generated from RAW264.7 cells using the High-Capacity cDNA Reverse Transcription Kit (Thermo Fisher; catalog # 4368814) and was subsequently cloned into p3x-FLAG-CMV10 mammalian expression vector. A catalytically inactive S172A mutant of mouse LIPA was generated in the same plasmid using a standard site-directed mutagenesis approach with Phusion polymerase & DpnI digestion (New England Biolabs) as per manufacturer’s instructions. The WT and S172A variants of LIPA were transfected in HEK293T cells using polyethyleneimine “max” 40,000 MW (Polysciences, Inc.) using a transient transfection strategy reported by us^46,56^. As established earlier, an empty vector “mock” was used in this experiment, to control for any off-target effects of the plasmid. Overexpression of WT and S172A variants of LIPA in HEK293T cells, relative to the mock control were confirmed by Western blot analysis. The primary antibodies used in this Western blot analysis were anti-Flag (mouse IgG) (Invitrogen; catalog # 14-6681-80) and anti-GAPDH (rabbit IgG) (Sigma-Aldrich; catalog # G9545) at a 1:1000 dilution. The secondary antibodies used in this Western blot analysis were HRP-conjugated anti-rabbit Ab (Goat IgG) (Invitrogen; catalog # 31460) and HRP-conjugated anti-mouse Ab (Goat IgG) (Abcam; catalog # ab6789) at a 1:10,000 dilution. All Western blot analysis were done on a PVDF membrane (GE Healthcare), developed by the Immobilon Western Chemiluminescent HRP substrate (Millipore; catalog # WBKLSO500) and imaged on a Syngene G-Box Chemi-XRQ gel documentation system. Gel-based ABPP and LC-MS based CE hydrolase analysis was performed as reported in earlier sections with 50 and 40 μg of protein lysate respectively in DPBS at pH 7.5, or 50 mM sodium acetate buffer at pH 4.5.

### Bead uptake and phagocytic index

To assess the uptake and intracellular localisation of beads, RAW264.7 cells were cultured on lysine coated coverslips up to 80% confluency. The cells were then treated with LAL at 20 μM final concentration for 4 hours, following which, they were fed with 200 µL solution of 1-µm fluorescent polystyrene beads (Polysciences; catalog # 15702) in DMEM. Thereafter, the cells were kept at 4 °C to synchronize the uptake of beads and then pulsed for 15 mins at 37 ^о^C. The cells were then washed with cold sterile DPBS (3x) and incubated at 37 ^о^C for another 4 hours. Post incubation, the cells were washed with cold sterile DPBS and fixed using 4% (w/v) paraformaldehyde (PFA) (Sigma-Aldrich; catalog # P6148) (for 15 min). To mark the nucleus, the cells were stained with 4’,6-diamidino-2-phenylindole (DAPI), washed with cold sterile DPBS (3x) and placed on a slide using a mounting media (Invitrogen; catalog # S36937). The fixed cells were imaged at ambient temperature on an inverted confocal microscope (LSM710 or LSM780 from Zeiss). Marking the periphery of cells, counting the number of beads phagocytosed per cell (**Figure 3A**), and measuring the intracellular distances of beads from the centre of the nucleus/cellular periphery (**Figure 3B**, **3C**) were all done using ImageJ software^60,61^.

### Immunofluorescence (IFA) assays

All IFA assays with phagosomes were done using protocols previously reported by us with some modifications^25^. Briefly, acid-washed coverslips were incubated for 20 min at ambient temperature in a poly-*L*-lysine solution (Sigma-Aldrich; catalog # P8920) [prepared by mixing 18 mL ethanol (analytical reagent grade, 99.9%) (MSD Chemicals, catalog # 5268-39), 2 mL MeOH (analytical reagent grade, 99.9%) (Fisher Chemicals, catalog # A456-4), and 1 mL poly-*L*-lysine (0.1% w/v in sterile MilliQ water)]. The coverslips were subsequently dried at 100 °C for 10 min, cooled to ambient temperature, and transferred to a 24-well plate. 20 μL EP or LP (vehicle and inhibitor treated) preparations (prepared using 3-µm silica beads) were added to the coverslips and centrifuged at 1200*g* for 5 min using a swinging bucket rotor (Eppendorf). Post-centrifuging, the coverslips were washed with sterile DPBS (1x), fixed with 4% (w/v) PFA (15 min at ambient temperature), and blocked with 6% (w/v) bovine serum albumin (Sigma-Aldrich; catalog # SRE0098) in DPBS (60 min at ambient temperature). The antibody incubation procedure was similar to that reported earlier^25^. The primary antibodies used in this study were: rabbit monoclonal anti-Flotillin-1 (Abcam; catalog # EPR6041), rabbit polyclonal anti-LAMP1 (Abcam; catalog # ab24170), rabbit polyclonal anti-EEA1 (Abcam; catalog # ab264245); all at a dilution of 1:250. The secondary antibody used in this study was the donkey anti-rabbit Alexa Fluor-488 conjugated antibody (Thermo Fisher Scientific; catalog # A-21206) at a dilution of 1:1000. All antibody labelled phagosomes were imaged on a LSM 510 confocal microscope (Zeiss). The outlining of the phagosome boundary, and the measurement of fluorescence intensity were done using the ImageJ software^60,61^. In all our experiments, the background intensities were also measured, and subtracted from the total fluorescence intensity obtained on phagosomes.

### Microbial clearance assays

**(A) *S. aureus***: The J774.A1 murine macrophage cell line was seeded at 50,000 cells/well in a 12-well tissue culture plate. After 18 hours, one set of cells were treated with 20 μM LAL and this concentration was maintained in the media throughout the course experiment, while other set of cells remained untreated (treated with vehicle, DMSO). After 4 hours of LAL (or vehicle) treatment, both group of cells were infected with *S. aureus* (ATCC, catalog # 29213) with a multiplicity of infection (MOI) at 1:100 and incubated at 37 °C with 5% (v/v) CO_2_ for 90 min. The inoculum was prepared in sterile DPBS and finally diluted in cell culture media to let the bacterial cells acclimatize before infection, and this inoculum was plated on Tryptic Soya Agar (TSA) plates to confirm the MOI. After the 90 min infection period, the cells were washed with sterile DBPS (3x) and treated with lysostaphin 10U (30 min) to remove the extracellular bacterial load. Thereafter, the cells were washed again with sterile DPBS (3x) and fresh media containing lysostaphin (4U) added. In each experiment, colony forming unit (CFU) plating were done before and after the lysostaphin treatment to ensure the eradication of extracellular bacterial load. At each experimental time point (4, 8 and 24 hour post infection), the cells from both the treatment groups (LAL and vehicle treated) were harvested, lysed using RIPA buffer, and the intracellular bacterial load was plated on TSA plates, that were kept at 37 °C for overnight. The next day, colonies were counted, and the log_10_[CFU/mL inoculum] was calculated. **(B) *M. tuberculosis***: Intracellular *M. tuberculosis* survival was determined by the drop-down CFU method^62^. RAW264.7 cells were cultured as mentioned earlier, seeded in a 96-well plate at 15000 cells/well, and treated with LAL (20 μM, 4 hours) or vehicle (DMSO). From this point on, 20 μM LAL concentration was maintained in the media throughout the course experiment, while other set of cells remained untreated (treated with vehicle, DMSO). The *M. tuberculosis* culture (of laboratory strain H37Rv) was grown to 0.4 - 0.6 (based on optical density (OD) at 600 nm), washed and passed through a 26-gauge needle and 30-gauge needle (5x each) to make a single cell suspension. Thereafter, the bacterial number was estimated based on OD at 600nm for infection and 10 MOI was added to LAL or vehicle treated RAW264.7 cells. After the 4 hour infection period, the cells were washed with sterile DMEM (2x) to remove extracellular bacteria. Fresh media with amikacin (100 mg/mL) was added to the infected RAW264.7 cells and incubated for 2 hours to eradicate any remaining extracellular bacteria. Following that, cells were washed with DMEM (2x), supplemented with fresh media containing LAL (20 μM), and cultured till the experimental time points (24, 48 and 72 hours). At each experimental time point, the cell culture media was removed, the cells were washed with sterile DPBS (2x), lysed with a solution 0.06% (w/v) sodium dodecyl sulfate for 10 min at room temperature. The required dilutions (N/10 and N/50) were prepared with 7H9 media in a fresh 96-well plate, and 10 μl of lysed cells dilutions were plated on a 7H11 agar plate by the drop-down method^62^. The plates were allowed to dry, and then incubated for 18 to 21 days at 37 °C, following which, individual bacterial colonies were counted manually and converted to CFU/mL, by accounting for the dilution factor and the fraction of volume used for CFU plating. In parallel, MTT assays were also performed to score for macrophage viability at each time point, where, instead of lysing the cells, MTT solution was added, and the OD was taken at 595 nm.

### Statistical analysis

All statistical analyses were performed using the GraphPad Prism 10 (version 10.6.0) for macOS software. All data are shown as mean ± standard deviation. The Student’s two-tailed *t*-test was used to ascertain statistically significant difference between the different experimental groups, and a *P-*value < 0.05 was considered statistically significant for this study.

## DATA AVAILABILITY

The data supporting the findings of this study are available in the manuscript, its Supplementary Information files, or are available with the corresponding author upon reasonable request. Additionally, publicly available datasets associated with this manuscript include: (1) <u>Proteomics data</u>: All the LC-MS based proteomics data is deposited in the PRIDE repository (accession number PXD051170). (2) <u>Untargeted lipidomics data</u>: The raw untargeted lipidomics data has been deposited in Mendeley.

## SUPPLEMENTARY INFORMATION

Supplementary Information includes Supplementary Figures and Supplementary Table 1.

## ETHICAL DECLARATIONS

The authors declare no competing interests.

## ACKNOWLEDGEMENTS

This study was financially supported by a SwarnaJayanti Fellowship, Science and Engineering Research Board (SERB), Department of Science and Technology, Government of India (grant number: SB/SJF/2021–22/01 to S.S.K.), EMBO Young Investigator’s Award (to S.S.K.), Department of Biotechnology (DBT) – Wellcome Trust India Alliance Fellowship (grant numbers: IA/S/11/2500255 and IA/S/19/2/504634 to R. M.), Prime Minister’s Research Fellowship (graduate student fellowship to O.S.) and DBT Fellowship (grant number: (DBT/21-22/IIT-B/1603 to B.). Additionally, intramural funding support from IISER Pune (to S.S.K.) and IIT Bombay (to R.M.) are also acknowledged. We thank Saddam Shekh for technical assistance and maintenance of the biological mass spectrometry facility at IISER Pune. Members of the S.S.K. lab at IISER Pune and R.M. lab at IIT Mumbai are thanked for providing critical comments and inputs throughout the course of this study. We acknowledge Tuberculosis Aerosol Challenge Facility, a DBT-supported BSL3 facility at ICGEB New Delhi, for conducting *M. tuberculosis* infection experiments.

## AUTHOR CONTRIBUTION

O.S. performed all the experiments; B. assisted O.S. with all the microscopy experiments under the supervision of R.M.; S.S.K.S. and D.K. performed the MTB experiments; A. A. and S.C. performed the S. aureus experiments; O.S., R. M. and S.S.K. conceived the project; S.S.K. supervised the project and acquired funding for the project; O.S. and S.S.K. wrote the manuscript with inputs from all authors.

## REFERENCES

1. King, R.J., Singh, P.K., and Mehla, K. (2022). The cholesterol pathway: impact on immunity and cancer. Trends Immunol 43, 78–92. 10.1016/j.it.2021.11.007.

2. Griffiths, W.J., and Wang, Y. (2022). Cholesterol metabolism: from lipidomics to immunology. J Lipid Res 63, 100165. 10.1016/j.jlr.2021.100165.

3. Rao, M., and Mayor, S. (2014). Active organization of membrane constituents in living cells. Curr Opin Cell Biol 29, 126–132. 10.1016/j.ceb.2014.05.007.

4. Mayor, S., and Rao, M. (2004). Rafts: scale-dependent, active lipid organization at the cell surface. Traffic 5, 231–240. 10.1111/j.1600-0854.2004.00172.x.

5. Ruysschaert, J.M., and Lonez, C. (2015). Role of lipid microdomains in TLR-mediated signalling. Biochim Biophys Acta 1848, 1860–1867. 10.1016/j.bbamem.2015.03.014.

6. de la Roche, M., Hamilton, C., Mortensen, R., Jeyaprakash, A.A., Ghosh, S., and Anand, P.K. (2018). Trafficking of cholesterol to the ER is required for NLRP3 inflammasome activation. J Cell Biol 217, 3560–3576. 10.1083/jcb.201709057.

7. Westerterp, M., Fotakis, P., Ouimet, M., Bochem, A.E., Zhang, H., Molusky, M.M., Wang, W., Abramowicz, S., la Bastide-van Gemert, S., Wang, N., et al. (2018). Cholesterol Efflux Pathways Suppress Inflammasome Activation, NETosis, and Atherogenesis. Circulation 138, 898–912. 10.1161/CIRCULATIONAHA.117.032636.

8. Galvan-Pena, S., and O’Neill, L.A. (2014). Metabolic reprograming in macrophage polarization. Front Immunol 5, 420. 10.3389/fimmu.2014.00420.

9. Liu, Y., Xu, R., Gu, H., Zhang, E., Qu, J., Cao, W., Huang, X., Yan, H., He, J., and Cai, Z. (2021). Metabolic reprogramming in macrophage responses. Biomark Res 9, 1. 10.1186/s40364-020-00251-y.

10. Duan, Y.J., Gong, K., Xu, S.W., Zhang, F., Meng, X.S., and Han, J.H. (2022). Regulation of cholesterol homeostasis in health and diseases: from mechanisms to targeted therapeutics. Signal Transduct Tar 7, 265. https://doi.org/ARTN 265 10.1038/s41392-022-01125-5.

11. Gui, Y., Zheng, H., and Cao, R.Y. (2022). Foam Cells in Atherosclerosis: Novel Insights Into Its Origins, Consequences, and Molecular Mechanisms. Front Cardiovasc Med 9, 845942. 10.3389/fcvm.2022.845942.

12. Guerrini, V., and Gennaro, M.L. (2019). Foam Cells: One Size Doesn’t Fit All. Trends Immunol 40, 1163–1179. 10.1016/j.it.2019.10.002.

13. Angelovich, T.A., Hearps, A.C., and Jaworowski, A. (2015). Inflammation-induced foam cell formation in chronic inflammatory disease. Immunol Cell Biol 93, 683–693. 10.1038/icb.2015.26.

14. Pieters, J. (2001). Entry and survival of pathogenic mycobacteria in macrophages. Microbes Infect 3, 249–255. 10.1016/s1286-4579(01)01376-4.

15. Ferrari, G., Langen, H., Naito, M., and Pieters, J. (1999). A coat protein on phagosomes involved in the intracellular survival of mycobacteria. Cell 97, 435–447. 10.1016/s0092-8674(00)80754-0.

16. Allen, L.A. (2007). Phagocytosis and persistence of Helicobacter pylori. Cell Microbiol 9, 817–828. 10.1111/j.1462-5822.2007.00906.x.

17. Ramarao, N., and Meyer, T.F. (2001). Helicobacter pylori resists phagocytosis by macrophages: quantitative assessment by confocal microscopy and fluorescence-activated cell sorting. Infect Immun 69, 2604–2611. 10.1128/IAI.69.4.2604-2611.2001.

18. Moldovan, A., and Fraunholz, M.J. (2019). In or out: Phagosomal escape of Staphylococcus aureus. Cell Microbiol 21, e12997. 10.1111/cmi.12997.

19. Thammavongsa, V., Kim, H.K., Missiakas, D., and Schneewind, O. (2015). Staphylococcal manipulation of host immune responses. Nat Rev Microbiol 13, 529–543. 10.1038/nrmicro3521.

20. Walpole, G.F.W., Grinstein, S., and Westman, J. (2018). The role of lipids in host-pathogen interactions. IUBMB Life 70, 384–392. 10.1002/iub.1737.

21. Goluszko, P., and Nowicki, B. (2005). Membrane cholesterol: a crucial molecule affecting interactions of microbial pathogens with mammalian cells. Infect Immun 73, 7791–7796. 10.1128/IAI.73.12.7791-7796.2005.

22. Levin, R., Grinstein, S., and Canton, J. (2016). The life cycle of phagosomes: formation, maturation, and resolution. Immunol Rev 273, 156–179. 10.1111/imr.12439.

23. Flannagan, R.S., Jaumouille, V., and Grinstein, S. (2012). The cell biology of phagocytosis. Annu Rev Pathol 7, 61–98. 10.1146/annurev-pathol-011811-132445.

24. Saharan, O., and Kamat, S.S. (2023). Mapping lipid pathways during phagocytosis. Biochem Soc Trans 51, 1279–1287. 10.1042/BST20221424.

25. Rai, A., Pathak, D., Thakur, S., Singh, S., Dubey, A.K., and Mallik, R. (2016). Dynein Clusters into Lipid Microdomains on Phagosomes to Drive Rapid Transport toward Lysosomes. Cell 164, 722–734. 10.1016/j.cell.2015.12.054.

26. Mehendale, N., Mallik, R., and Kamat, S.S. (2021). Mapping Sphingolipid Metabolism Pathways during Phagosomal Maturation. ACS Chem Biol 16, 2757–2765. 10.1021/acschembio.1c00393.

27. Lebrand, C., Corti, M., Goodson, H., Cosson, P., Cavalli, V., Mayran, N., Faure, J., and Gruenberg, J. (2002). Late endosome motility depends on lipids via the small GTPase Rab7. EMBO J 21, 1289–1300. 10.1093/emboj/21.6.1289.

28. Huynh, K.K., Gershenzon, E., and Grinstein, S. (2008). Cholesterol accumulation by macrophages impairs phagosome maturation. J Biol Chem 283, 35745–35755. 10.1074/jbc.M806232200.

29. Chandramouli, A., and Kamat, S.S. (2024). A Facile LC-MS Method for Profiling Cholesterol and Cholesteryl Esters in Mammalian Cells and Tissues. Biochemistry 63, 2300–2309. 10.1021/acs.biochem.4c00160.

30. Pathak, D., Mehendale, N., Singh, S., Mallik, R., and Kamat, S.S. (2018). Lipidomics Suggests a New Role for Ceramide Synthase in Phagocytosis. ACS Chem Biol 13, 2280– 2287. 10.1021/acschembio.8b00438.

31. Kinchen, J.M., and Ravichandran, K.S. (2008). Phagosome maturation: going through the acid test. Nat Rev Mol Cell Biol 9, 781–795. 10.1038/nrm2515.

32. Long, J.Z., and Cravatt, B.F. (2011). The metabolic serine hydrolases and their functions in mammalian physiology and disease. Chem Rev 111, 6022–6063.

33. Niphakis, M.J., and Cravatt, B.F. (2014). Enzyme inhibitor discovery by activity-based protein profiling. Annu Rev Biochem 83, 341–377. 10.1146/annurev-biochem-060713-035708.

34. Shui, W.Q., Sheu, L., Liu, J., Smart, B., Petzold, C.J., Hsieh, T.Y., Pitcher, A., Keasling, J.D., and Bertozzi, C.R. (2008). Membrane proteomics of phagosomes suggests a connection to autophagy. P Natl Acad Sci USA 105, 16952–16957. 10.1073/pnas.0809218105.

35. Rogers, L.D., and Foster, L.J. (2007). The dynamic phagosomal proteome and the contribution of the endoplasmic reticulum. Proc Natl Acad Sci U S A 104, 18520–18525. 10.1073/pnas.0705801104.

36. Li, F., and Zhang, H. (2019). Lysosomal Acid Lipase in Lipid Metabolism and Beyond. Arterioscler Thromb Vasc Biol 39, 850–856. 10.1161/ATVBAHA.119.312136.

37. Zhang, H. (2018). Lysosomal acid lipase and lipid metabolism: new mechanisms, new questions, and new therapies. Curr Opin Lipidol 29, 218–223. 10.1097/MOL.0000000000000507.

38. Ameis, D., Merkel, M., Eckerskorn, C., and Greten, H. (1994). Purification, characterization and molecular cloning of human hepatic lysosomal acid lipase. Eur J Biochem 219, 905–914. 10.1111/j.1432-1033.1994.tb18572.x.

39. Warner, T.G., Dambach, L.M., Shin, J.H., and O’Brien, J.S. (1981). Purification of the lysosomal acid lipase from human liver and its role in lysosomal lipid hydrolysis. J Biol Chem 256, 2952–2957.

40. Bradic, I., Kuentzel, K.B., Honeder, S., Grabner, G.F., Vujic, N., Zimmermann, R., Birner-Gruenberger, R., and Kratky, D. (2022). Off-target effects of the lysosomal acid lipase inhibitors Lalistat-1 and Lalistat-2 on neutral lipid hydrolases. Mol Metab 61, 101510. 10.1016/j.molmet.2022.101510.

41. Hamilton, J., Jones, I., Srivastava, R., and Galloway, P. (2012). A new method for the measurement of lysosomal acid lipase in dried blood spots using the inhibitor Lalistat 2. Clin Chim Acta 413, 1207–1210. 10.1016/j.cca.2012.03.019.

42. Chang, J.W., Nomura, D.K., and Cravatt, B.F. (2011). A potent and selective inhibitor of KIAA1363/AADACL1 that impairs prostate cancer pathogenesis. Chem Biol 18, 476–484. 10.1016/j.chembiol.2011.02.008.

43. Kinsey, S.G., Long, J.Z., O’Neal, S.T., Abdullah, R.A., Poklis, J.L., Boger, D.L., Cravatt, B.F., and Lichtman, A.H. (2009). Blockade of endocannabinoid-degrading enzymes attenuates neuropathic pain. J Pharmacol Exp Ther 330, 902–910. 10.1124/jpet.109.155465.

44. Hsu, K.L., Tsuboi, K., Adibekian, A., Pugh, H., Masuda, K., and Cravatt, B.F. (2012). DAGLbeta inhibition perturbs a lipid network involved in macrophage inflammatory responses. Nat Chem Biol. 8, 999–1007.

45. Mayer, N., Schweiger, M., Romauch, M., Grabner, G.F., Eichmann, T.O., Fuchs, E., Ivkovic, J., Heier, C., Mrak, I., Lass, A. et al. (2013). Development of small-molecule inhibitors targeting adipose triglyceride lipase. Nat Chem Biol 9, 785–787. 10.1038/nchembio.1359.

46. Kelkar, D.S., Ravikumar, G., Mehendale, N., Singh, S., Joshi, A., Sharma, A.K., Mhetre, A., Rajendran, A., Chakrapani, H., and Kamat, S.S. (2019). A chemical-genetic screen identifies ABHD12 as an oxidized-phosphatidylserine lipase. Nat Chem Biol 15, 169–178. 10.1038/s41589-018-0195-0.

47. Garin, J., Diez, R., Kieffer, S., Dermine, J.F., Duclos, S., Gagnon, E., Sadoul, R., Rondeau, C., and Desjardins, M. (2001). The phagosome proteome: insight into phagosome functions. J Cell Biol 152, 165–180. 10.1083/jcb.152.1.165.

48. Chambers, H.F., and Deleo, F.R. (2009). Waves of resistance: Staphylococcus aureus in the antibiotic era. Nat Rev Microbiol 7, 629–641. 10.1038/nrmicro2200.

49. Lowy, F.D. (2003). Antimicrobial resistance: the example of Staphylococcus aureus. J Clin Invest 111, 1265–1273. 10.1172/JCI18535.

50. Trajman, A., Campbell, J.R., Kunor, T., Ruslami, R., Amanullah, F., Behr, M.A., and Menzies, D. (2025). Tuberculosis. Lancet 405, 850–866. 10.1016/S0140-6736(24)02479-6.

51. Pandey, A.K., and Sassetti, C.M. (2008). Mycobacterial persistence requires the utilization of host cholesterol. Proc Natl Acad Sci U S A 105, 4376–4380. 10.1073/pnas.0711159105.

52. Desjardins, M. (1995). Biogenesis of phagolysosomes: the ‘kiss and run’ hypothesis. Trends Cell Biol 5, 183–186. 10.1016/s0962-8924(00)88989-8.

53. Asna Ashari, K., Azari-Yam, A., Shahrooei, M., and Ziaee, V. (2023). Wolman disease presenting with hemophagocytic lymphohistiocytosis syndrome and a novel LIPA gene variant: a case report and review of the literature. J Med Case Rep 17, 369. 10.1186/s13256-023-04116-4.

54. Alabbas, F., Elyamany, G., Alanzi, T., Ali, T.B., Albatniji, F., and Alfaraidi, H. (2021). Wolman’s disease presenting with secondary hemophagocytic lymphohistiocytosis: a case report from Saudi Arabia and literature review. BMC Pediatr 21, 72. 10.1186/s12887-021-02541-2.

55. Taurisano, R., Maiorana, A., De Benedetti, F., Dionisi-Vici, C., Boldrini, R., and Deodato, F. (2014). Wolman disease associated with hemophagocytic lymphohistiocytosis: attempts for an explanation. Eur J Pediatr 173, 1391–1394. 10.1007/s00431-014-2338-y.

56. Chakraborty, A., Punnamraju, P., Sajeevan, T., Kaur, A., Kolthur-Seetharam, U., and Kamat, S.S. (2025). Identification of ABHD6 as a lysophosphatidylserine lipase in the mammalian liver and kidneys. J Biol Chem 301, 108157. 10.1016/j.jbc.2025.108157.

57. Kumar, K., Mhetre, A., Ratnaparkhi, G.S., and Kamat, S.S. (2021). A Superfamily-wide Activity Atlas of Serine Hydrolases in Drosophila melanogaster. Biochemistry 60, 1312–1324. 10.1021/acs.biochem.1c00171.

58. Kumar, K., Pazare, M., Ratnaparkhi, G.S., and Kamat, S.S. (2024). CG17192 is a Phospholipase That Regulates Signaling Lipids in the Gut upon Infection. Biochemistry 63, 3000–3010. 10.1021/acs.biochem.4c00579.

59. Shanbhag, K., Mhetre, A.B., Saharan, O., Devarajan, A., Rai, A., Madhusudhan, M.S., Chakrapani, H., and Kamat, S.S. (2025). Chemoproteomics identifies protein ligands for monoacylglycerol lipids. Commun Chem 8, 197. 10.1038/s42004-025-01589-w.

60. Rueden, C.T., Schindelin, J., Hiner, M.C., DeZonia, B.E., Walter, A.E., Arena, E.T., and Eliceiri, K.W. (2017). ImageJ2: ImageJ for the next generation of scientific image data. BMC Bioinformatics 18, 529. 10.1186/s12859-017-1934-z.

61. Arena, E.T., Rueden, C.T., Hiner, M.C., Wang, S., Yuan, M., and Eliceiri, K.W. (2017). Quantitating the cell: turning images into numbers with ImageJ. Wiley Interdiscip Rev Dev Biol 6. 10.1002/wdev.260.

62. Kumar, D., Nath, L., Kamal, M.A., Varshney, A., Jain, A., Singh, S., and Rao, K.V. (2010). Genome-wide analysis of the host intracellular network that regulates survival of Mycobacterium tuberculosis. Cell 140, 731–743. 10.1016/j.cell.2010.02.012.

