## Supplementary Figures for "Lysosomal acid lipase regulates cholesterol metabolism during phagosomal maturation"

### SUPPLEMENTARY INFORMATION

Supplementary Figures: 1 – 12

Supplementary Table 1: See excel sheet

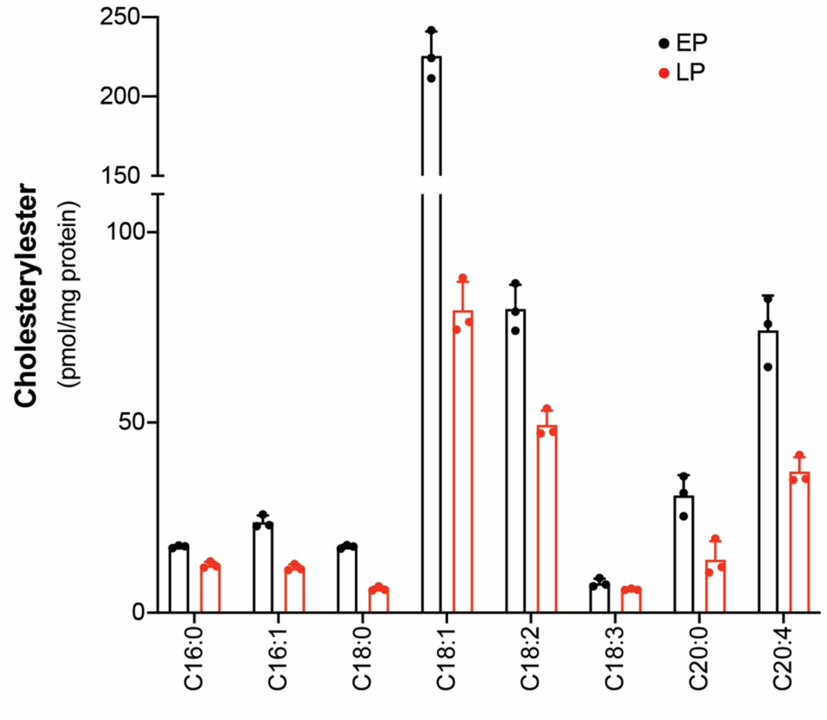


**Supplementary Figure 1.** Concentration of all the CEs on purified EPs (in black) and LPs (in red) measured by LC-MS analysis. The data represents mean ± standard deviation from three biological replicates per experimental group. See **Supplementary Table 1** for complete statistical analysis.

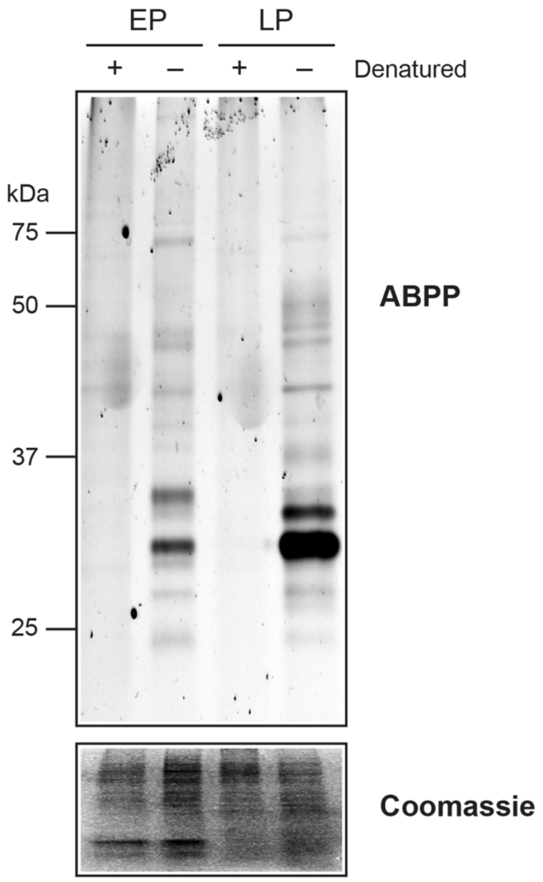


**Supplementary Figure 2.** A representative ABPP gel, showing activity of mSH profile of native or denatured lysates obtained from purified EPs or LPs. The bottom panel shows a Coomassie staining of the ABPP gel to confirm equal loading of lysates. This experiment was done three times with reproducible results each time.

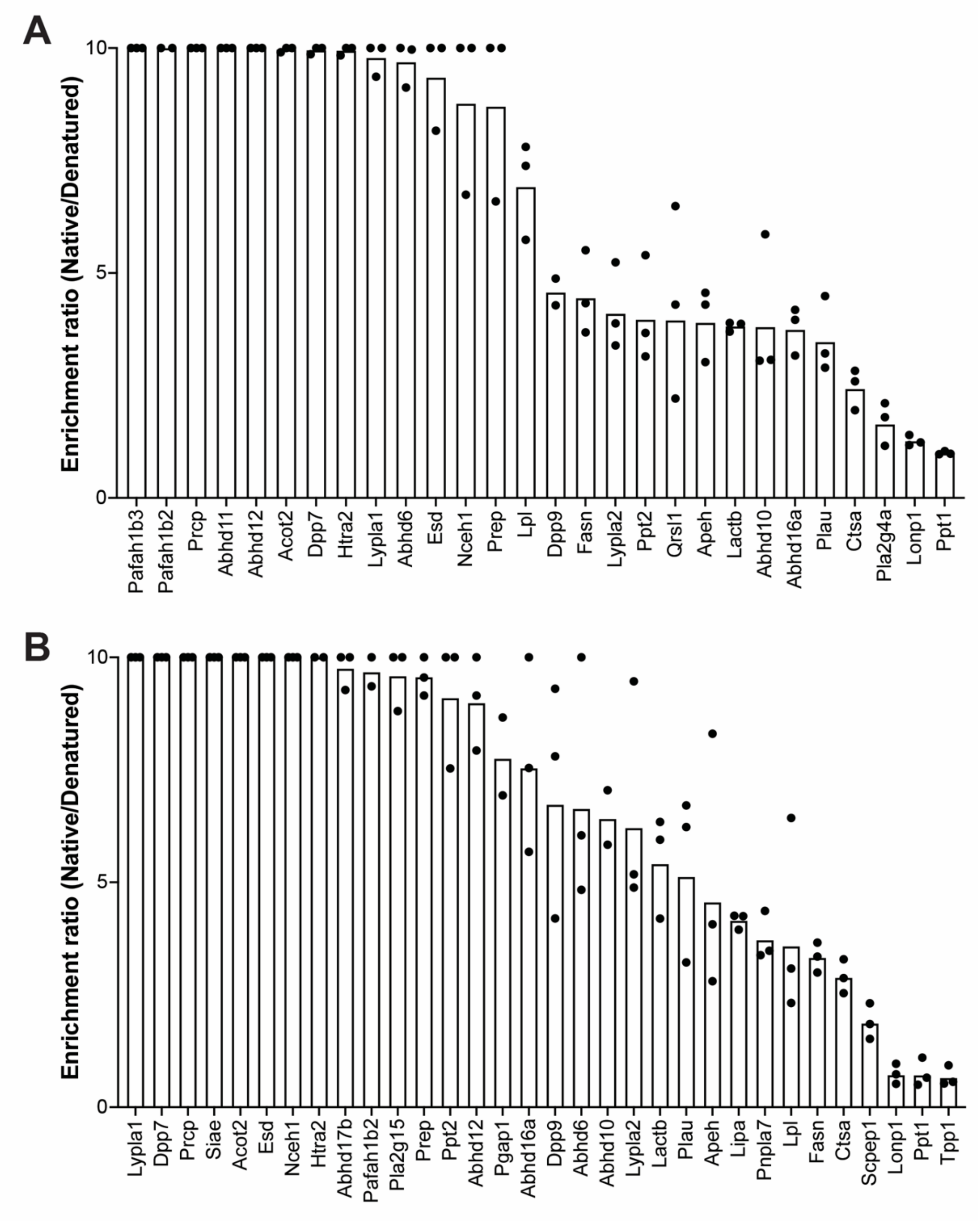


**Supplementary Figure 3. (A, B)** LC-MS based ABPP analysis quantitatively comparing mSH activities between native and denatured proteomes prepared from (**A**) purified EPs (**A**) or (**B**) purified LPs. The y-axis represents the enrichment ratio (native/denatured) for a particular mSH in this experiment. For (**A**), (**B**), data represents mean ± standard deviation from three biological replicates per experimental group. See **Supplementary Table 1** for complete details of enrichment ratios per replicate.

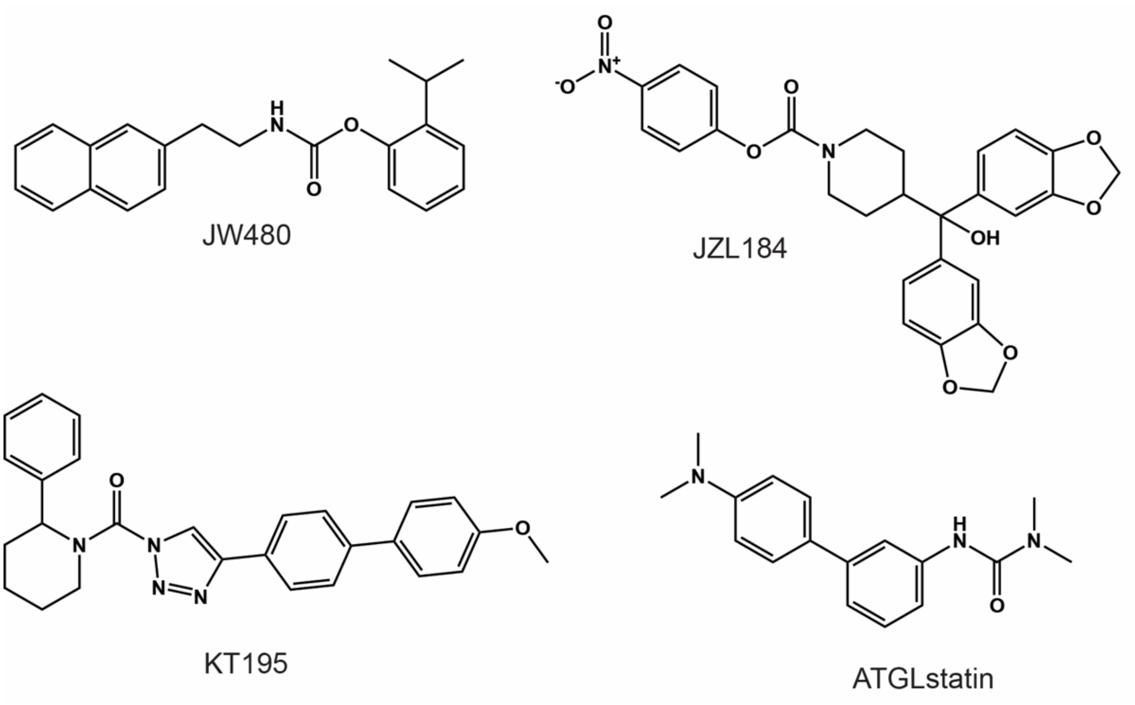


**Supplementary Figure 4.** Chemical structures of various specific inhibitors used in this study. The enzyme targets for these inhibitors are written in parenthesis: JW480 (NCEH1); JZL184 (MAGL); KT195 (ABHD6); and Atglistatin (ATGL).

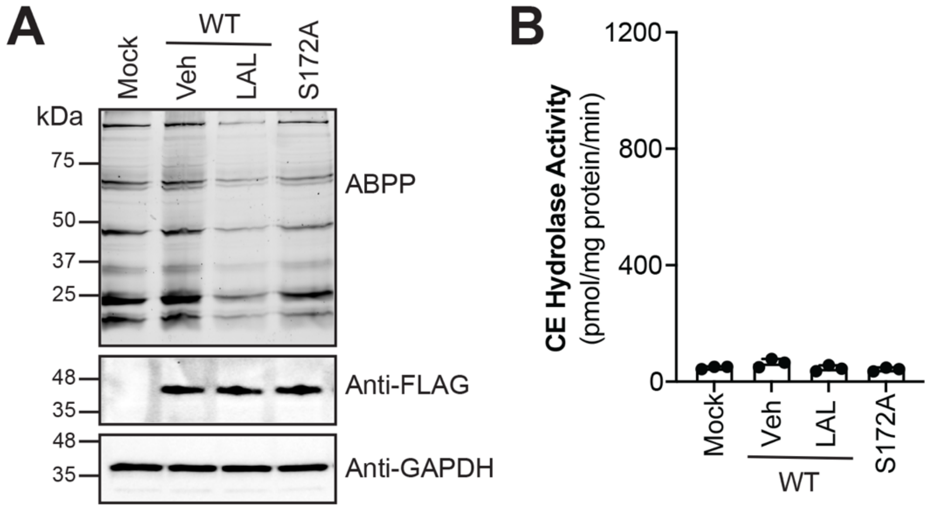


**Supplementary Figure 5.** (**A**) Lysates from HEK293T cells transfected with mock (empty vector), WT mouse LIPA (vehicle (DMSO) or LAL treated (20 μM, 30 min, 37 °C)), or the S172A variant of mouse LIPA were assessed using gel-based ABPP (top panel) (to check activity of WT and S172A LIPA in HEK293T cell lysates), Western blot analysis using an anti-FLAG antibody (middle panel) (to check overexpression of WT and S172A LIPA in HEK293T cell lysates), and an anti-GAPDH antibody (bottom panel) (to confirm equal protein loading). This experiment was done 3 times with reproducible results each time. (**B**) CE hydrolase activity of HEK293T cell lysates transfected with mock (empty vector), WT mouse LIPA (vehicle (DMSO) or LAL treated (20 μM, 30 min, 37 °C)), or the S172A variant of mouse LIPA. All the assays for (**A**) and (**B**) were done at pH 7.5. The data shown for (**B**) is plotted on the same scale as **Figure 2G** for comparative analysis. For (**B**) data represents mean ± standard deviation from three biological replicates per experimental group. See **Supplementary Table 1** for complete statistical analysis.

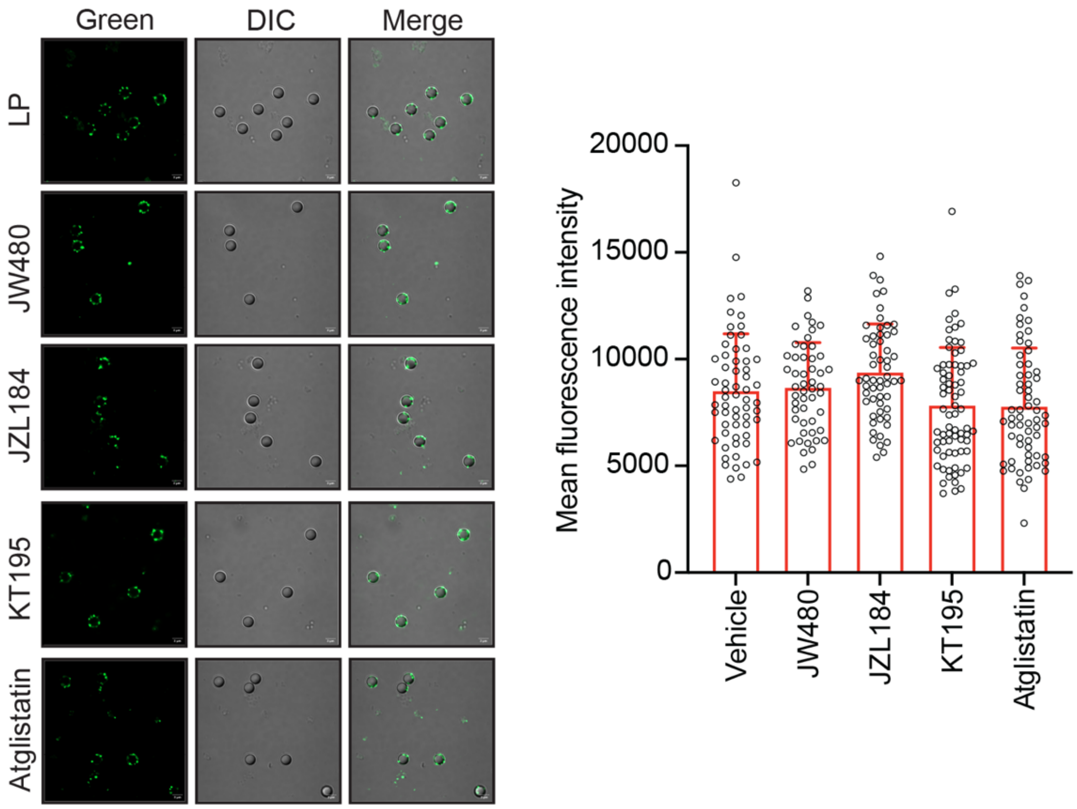


**Supplementary Figure 6.** *Left*, Representative images from the immunofluorescence analysis of purified LPs and inhibitor-treated LPs, using an anti-Flotilin-1 antibody (cholesterol-rich lipid raft marker). *Right*, Fluorescence intensity of individual phagosomes measured from the respective immunofluorescence experiment. The three images per experimental panel are a green immunofluorescence channel (Green) image (for a specific protein marker), a differential interference contrast (DIC) image (for outlining the phagosome), and an overlay of the Green and DIC images (Merge). The white line on all the images is a 3-μm scale. Data represents mean ± standard deviation from quantification of fluorescence intensity of >50 individual purified phagosomes per experimental group. All experiments were done at least three individual times with reproducible results each time. See **Supplementary Table 1** for complete statistical analysis for various experiments described in this figure.

| S.No. | Compound code | Solubility | MIC (µg/mL) | | | | | |
|---|---|---|---|---|---|---|---|---|
| | | | *E. coli* | *S. aureus* | *K. pneumoniae* | *A. baumannii* | *P. aeruginosa* | *M. tuberculosis* |
| 1 | Lalistat 2 | DMSO | >64 | >64 | >64 | >64 | >64 | >64 |
| 2 | Levofloxacin | DMSO | 0.0156 | 0.25 | 64 | 8 | 1 | 0.5 |

**Supplementary Figure 7.** Experimentally measured MIC values of LAL and a known antibiotic levofloxacin against a panel of pathogenic bacteria. The 64 µg/mL concentration of LAL corresponds to a molar concentration of > 200 µM, showing that LAL does not have bactericidal activity against any bacteria (specifically *S. aureus* and *M. tuberculosis*) at the 20 µM concentration used in this study.

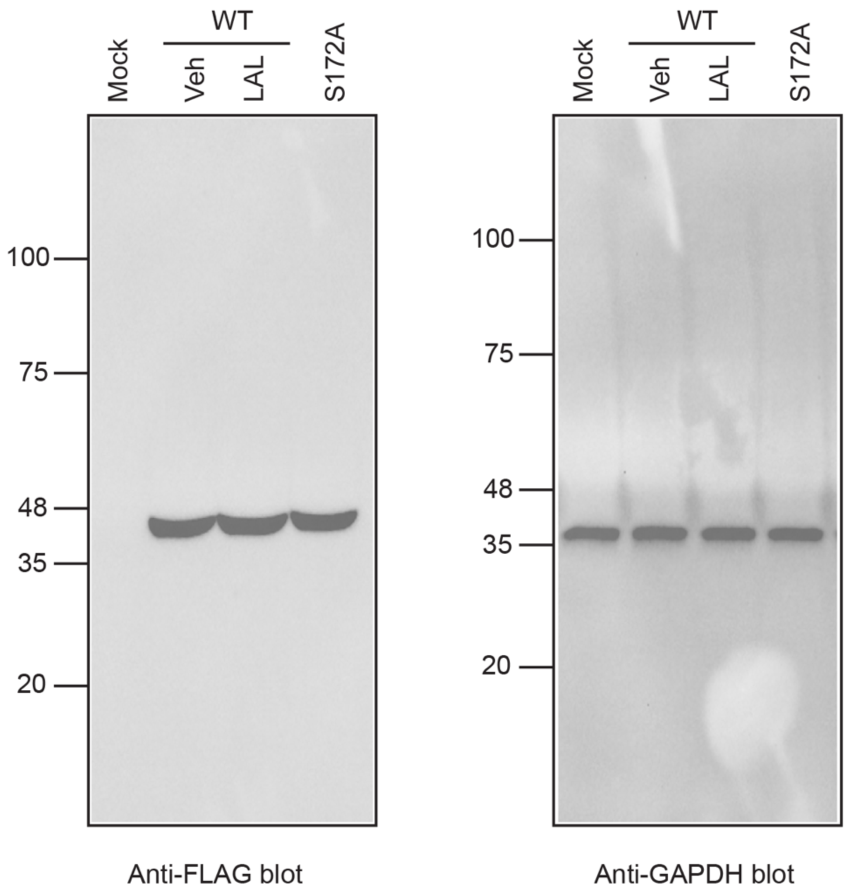


**Supplementary Figure 8.** Full blot images for Figure 2F.

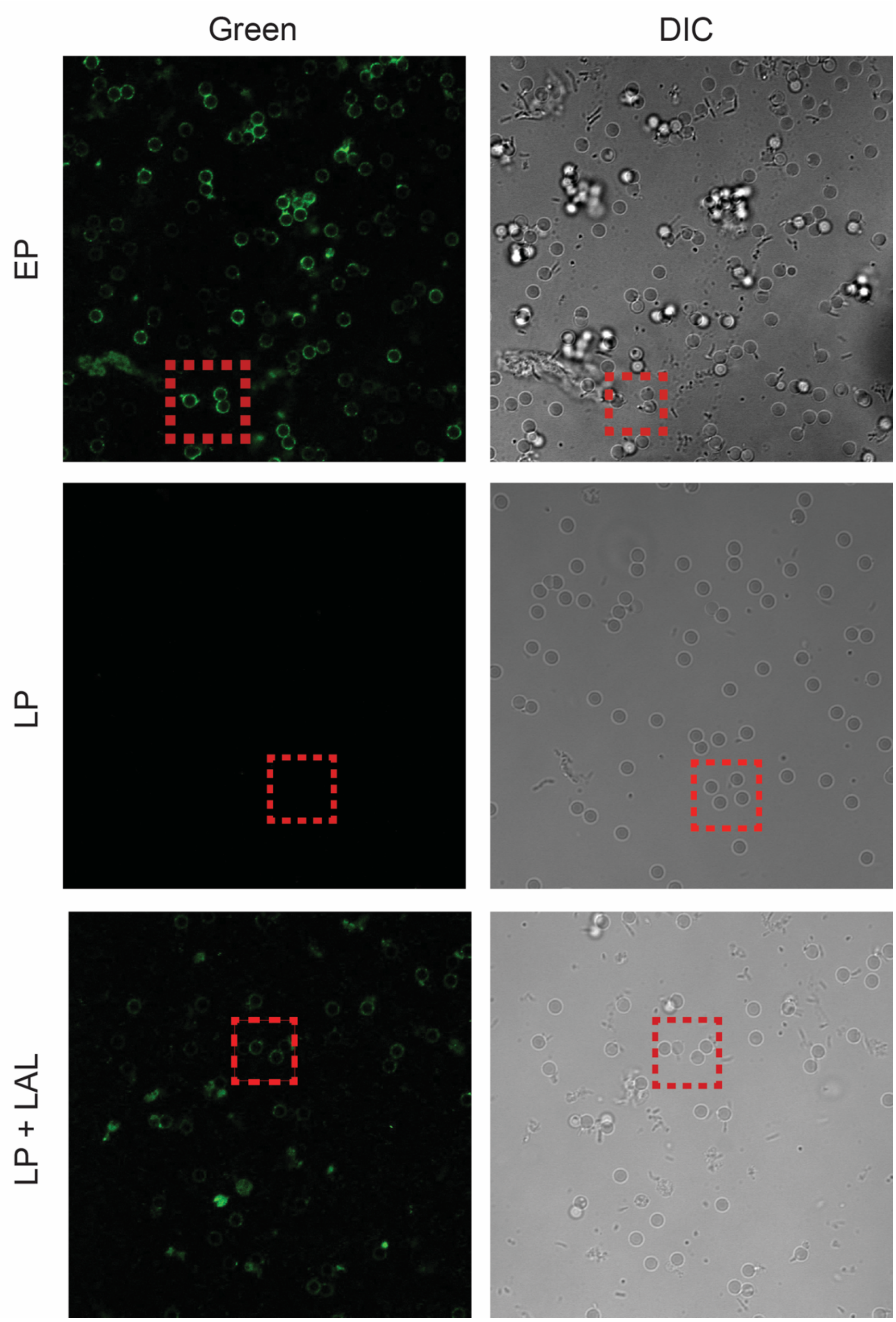


**Supplementary Figure 9.** Complete image for Figure 3D. Red box on each respective image shows zoomed part of whole image shown in Figure 3D.

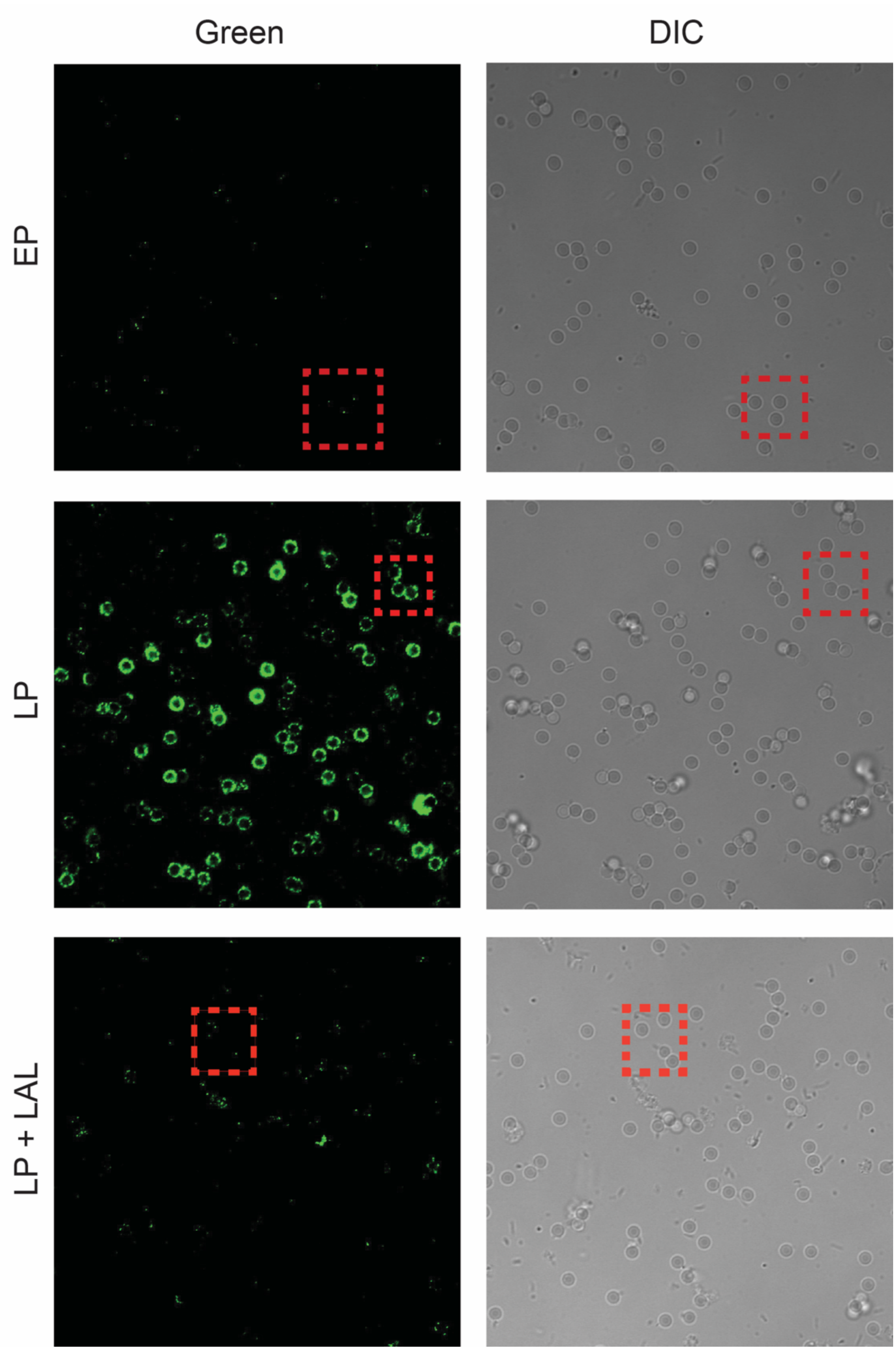


**Supplementary Figure 10.** Complete image for Figure 3E. Red box on each respective image shows zoomed part of whole image shown in Figure 3E.

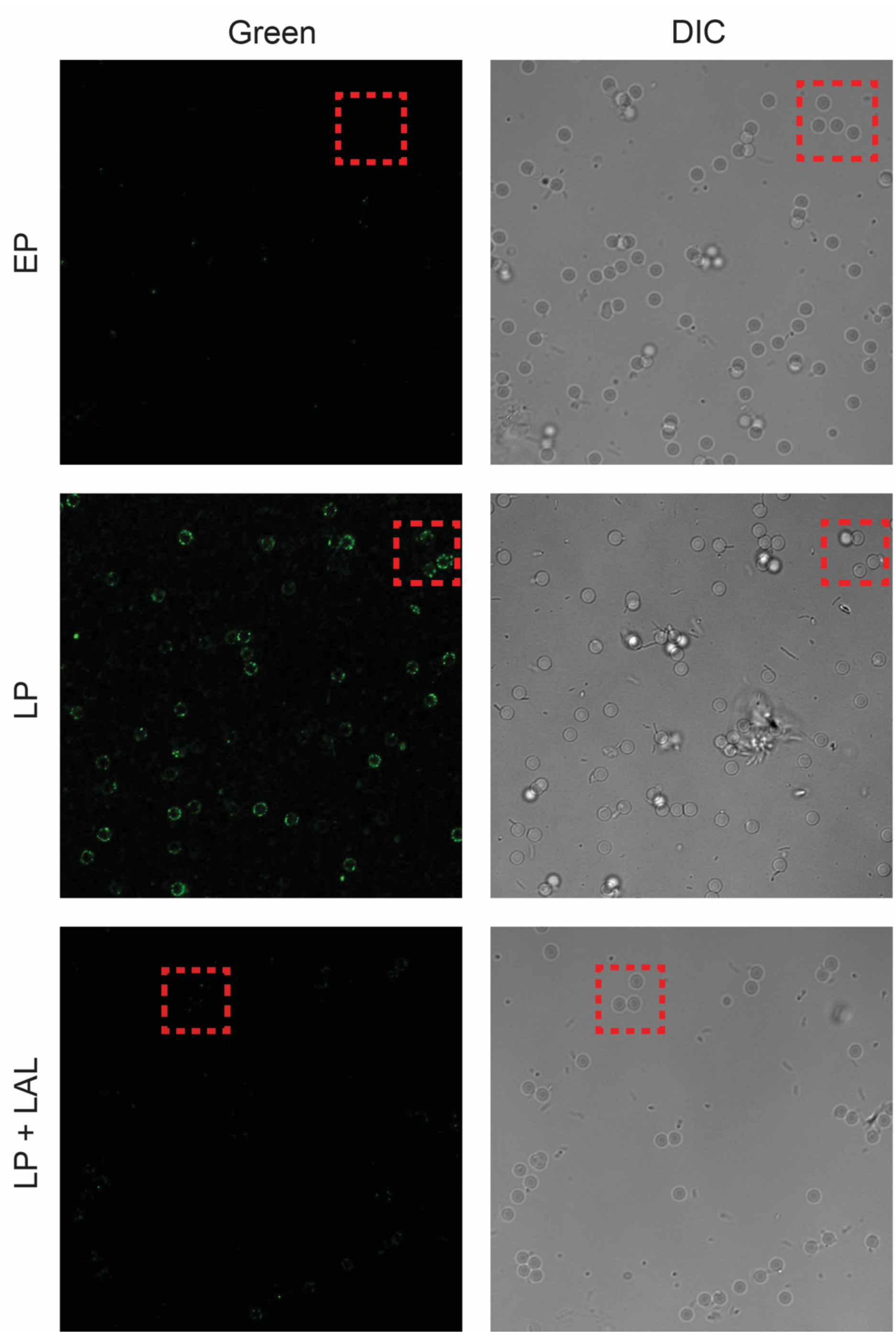


**Supplementary Figure 11.** Complete image for Figure 3F. Red box on each respective image shows zoomed part of whole image shown in Figure 3F.

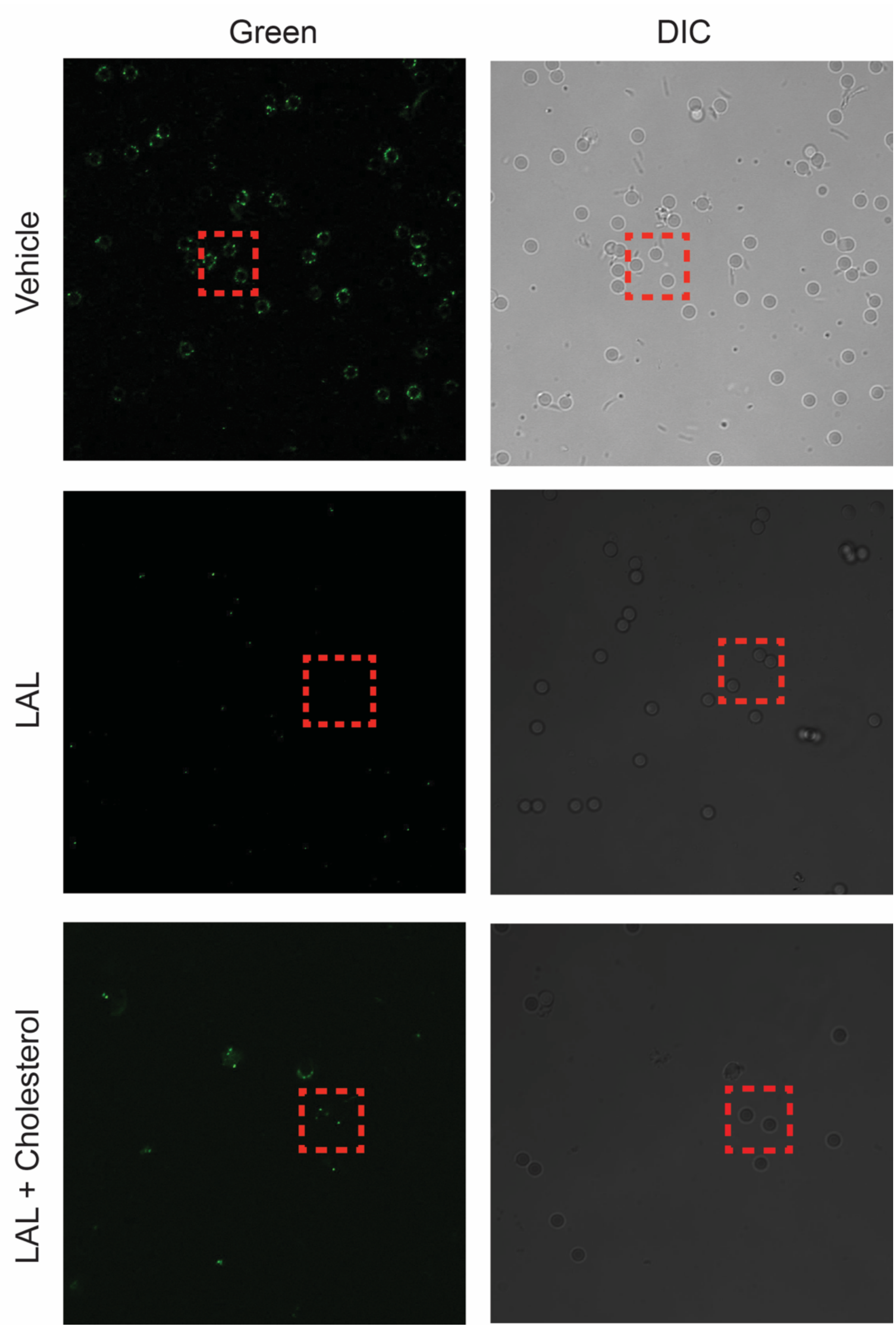


**Supplementary Figure 12.** Complete image for Figure 3G. Red box on each respective image shows zoomed part of whole image shown in Figure 3G.
